# NETISCE: A Network-Based Tool for Cell Fate Reprogramming

**DOI:** 10.1101/2021.12.30.474582

**Authors:** Lauren Marazzi, Milan Shah, Shreedula Balakrishnan, Ananya Patil, Paola Vera-Licona

## Abstract

The search for effective therapeutic targets in fields like regenerative medicine and cancer research has generated interest in cell fate reprogramming. This cellular reprogramming paradigm can drive cells to a desired target state from any initial state. However, methods for identifying reprogramming targets remain limited for biological systems that lack large sets of experimental data or a dynamical characterization. We present NETISCE, a novel computational tool for identifying cell fate reprogramming targets in static networks. In combination with machine learning algorithms, NETISCE estimates the attractor landscape and predicts reprogramming targets using Signal Flow Analysis and Feedback Vertex Set Control, respectively. Through validations in studies of cell fate reprogramming from developmental, stem cell, and cancer biology, we show that NETISCE can predict previously identified cell fate reprogramming targets and identify potentially novel combinations of targets. NETISCE extends cell fate reprogramming studies to larger-scale biological networks without the need for full model parameterization and can be implemented by experimental and computational biologists to identify parts of a biological system relevant to the desired reprogramming task.

## INTRODUCTION

Cell reprogramming redefines a cell’s identity by altering its epigenetic or transcriptional landscapes through the forced expression of transcription factors, small molecules, non-coding RNAs, or microenvironment-mediated changes. One type of cellular reprogramming, cell fate reprogramming, aims to control the internal state of a cell so that it is driven from a selected state to a target state or phenotype^1–7^. Practical applications of cell reprogramming in stem cell engineering^8–10^ and cancer biology^11–13^ have generated a great interest in the task of cell-fate reprogramming. Identifying combinations of reprogramming targets is especially useful for treating complex diseases, regenerating tissue, or reversing acquired resistance to canonical treatment regimens, where multi-drug approaches may be more effective than single drug therapy^1^,^14–18^.

Genome-wide and computational systems biology approaches are being broadly adopted for cell fate reprogramming studies. The majority of methods can be divided between those that take a network-based approach^19–24^ and those that take a dynamical systems-based approach^25–30^ to cell fate reprogramming^30–37^. However, these methods either do not capture all the essential information for cellular reprogramming^38^, or require mechanistic details and kinetic parameters to build a mathematical model of the system (for a brief overview of these approaches, please see Supplementary Text 1 and Supplementary Figure 1).

Although many network-based and dynamical systems-based methods take a trial-and-error approach to identify reprogramming targets, cell fate reprogramming can be viewed as a classical attractor-based control theory problem. The goal is to systematically determine how to shift the cell’s system from one attractor (stable state) to another with some degree of optimality. Previous theoretical studies on the controllability of systems show that even for large, non-linear biological systems, few targets need to be controlled to guide a system towards a biologically admissible target state^3,5^. This has been shown experimentally in cell fate reprogramming studies, such as in the transition of embryonic stem cells to somatic cells via knockdown of pluripotency-associated transcription factors^39^, the reversion of tumorigenesis by impairment of oncogenic signaling^40^, and fibroblast cell reprogramming^41^. The use of control theory to identify cellular reprogramming targets has been limited and is not directly applicable to large cell signaling networks^42,43^. Among the reasons for this limitation is the scarcity of available mechanistic details and kinetic parameters to build a mathematical model of the system, and, when the mechanistic rules are known, linear functions are used to describe them; however, it is unclear how the commonly observed switch-like behavior of biochemical processes^44,45^, can influence the results^1,27,46^.

A new method for identifying attractor-based reprogramming targets using only network topology extends from Control Theory for non-linear dynamics. The Feedback Vertex Set (FVS) Control is a structure-based, attractor-based control method suited to non-linear dynamical systems^5^. A network’s minimal FVS is the minimal set of nodes that intersects all cycles in a graph. FVS control states that appropriate perturbations on an FVS, which we refer to herein as FVS control nodes, can drive the system from any arbitrary initial state to any of the attractors of that system (Fig 1).

**Fig.1.**
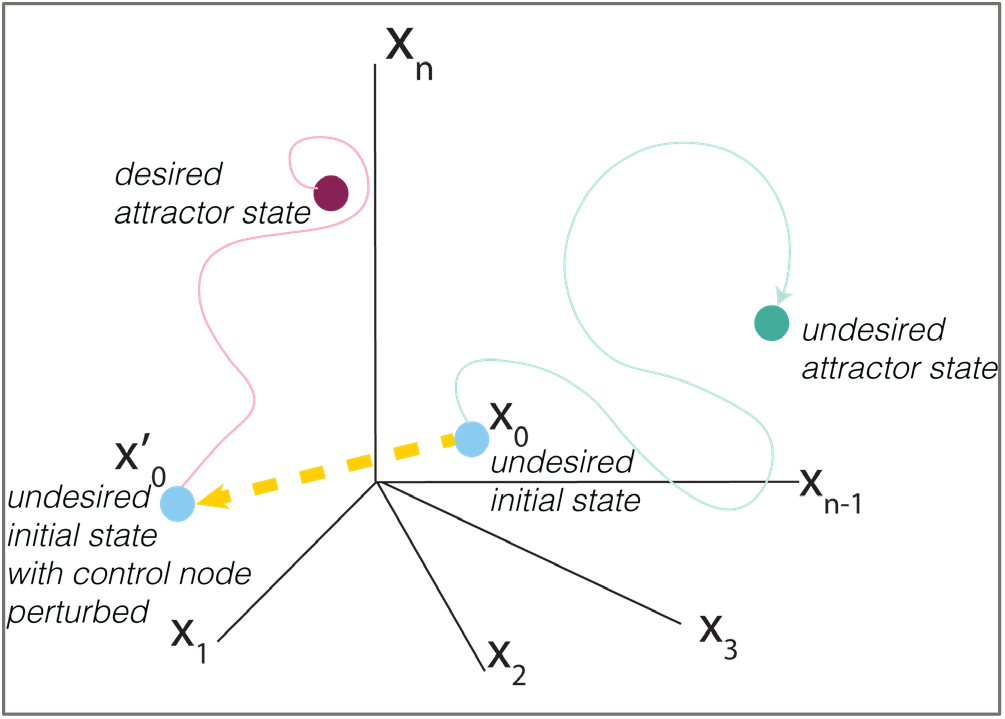
Control Theory View of Cell Fate Reprogramming. The Feedback Vertex Set control is a structure-based control method that can be applied to cell-fate reprogramming. By performing appropriate concerted perturbations to the minimal Feedback Vertex set on an initial state of the system that leads to an undesired attractor (green circle), the system can be shifted (yellow dashed lines) to a trajectory that leads to a desired attractor (magenta circle).

While the FVS control method provides a powerful cell fate reprogramming framework, it does not identify the specific perturbations needed on FVS control nodes (knockouts or overexpressions) to drive the system towards a particular set of attractors. By proceeding analogously to Boolean networks, estimating the system’s attractor landscape can aid in the search for the perturbations needed to be applied on FVS control nodes. To that end, signal flow estimation algorithms aim to estimate steady states without complete dynamical information of the system. They can be helpful to evaluate the effect of node perturbations in static networks^47,48^. The Signal Flow Analysis (SFA) method is especially suited to estimate system dynamics for non-linear complex systems^47^, and its application to biology has been recently explored^47,49^. SFA estimates a steady-state value for each network node based on a signal propagation equation that considers the activity of its regulators, the type of regulatory relationship (activation or inhibition), and the initial state of the node (see Methods). The signal propagation equation is solved for all network nodes synchronously until the difference between *x*(*t* + 1) and *x*(*t*) is less than a tolerance threshold. Thus, for a sufficiently small tolerance threshold (by default, this tolerance threshold is 10^-6^), a fixed-point attractor is generated for every initial state provided to SFA. As a result, unlike Boolean networks, where many initial states can arrive at one attractor, the attractor landscape estimated by SFA lacks large basins of attraction. In NETSICE, we use machine learning clustering algorithms to systematically partition the estimated attractors^50^ and produce the SFA-estimated attractor landscape. A phenotype is defined by the resultant set of attractors within a cluster and associated with the experimental sample whose attractor is contained in that cluster.

SFA has been used to successfully reproduce the steady-states of signaling networks derived from ODE models and the changes in expression for network elements under different perturbations from perturbation biology experiments with up to 80% accuracy^47^. Latterly, SFA was applied to an aging-related gene regulatory network (GRN) to identify potential aging reversion targets^49^. Although Lee and colleagues did not take a control theory-based approach, aging reversion targets were predicted by evaluating SFA-simulated single node perturbations that decreased the estimated steady-state expression values of aging-related biomarker nodes compared to an unperturbed simulation.

Previously, we introduced OCSANA+ (Optimal Control and Simulation of Signaling Networks from Network Analysis). This Cytoscape application implements an FVS finding algorithm and SFA to perform perturbation analyses on static networks^51^. OCSANA+ can be used to observe the effect of perturbations on FVS control nodes for biological networks in a user-friendly graphical interface. However, the application cannot customize the SFA algorithm to include gene expression data or perform an attractor landscape estimation. Additionally, the current software requires the user to configure every perturbation simulation manually.

In this work, we introduce NETISCE (NETwork-drIven analysiS of CEllular reprogramming), a novel computational method for identifying cell fate reprogramming targets. Specifically, in our context, we define cell fate reprogramming as guiding the system from an initial state to any of its attractors, which can be associated to an observable cell fate^43^. Our approach can be applied to any static network and only requires gene expression data from the undesired phenotype for the initial state of the system. NETISCE employs SFA and a machine learning clustering method to estimate the attractor landscape. Then SFA computes the attractor state reached by a given set of perturbations on an FVS control set. Finally, NETISCE uses machine learning classification methods to evaluate whether this newly computed attractor drives the system to the region of the attractor landscape associated with the desired cell fate.

To illustrate and validate our approach, we apply NETISCE to three different examples of cell fate reprogramming in developmental, stem cell, and cancer biology using GRNs and signaling networks. We show that NETISCE can reproduce the results of experimentally validated cell fate reprogramming studies and identify new reprogramming targets and perturbations.

We conclude that NETISCE extends the usefulness of static biological networks to analyses that currently require full parameterization. This is implemented by approaching cell fate reprogramming from the framework of control theory and dynamical systems and applying these concepts to a network-based analysis. Our method provides a practical and informative step for researchers designing experimental or mathematical modeling studies of cell fate reprogramming by identifying parts of the system relevant to the desired reprogramming task. NETISCE is a user-friendly tool implemented as a command-line Nextflow pipeline and Galaxy Project workflow that non-experts can use in modeling or computational approaches to analyze the biological systems of their interest.

## RESULTS

NETISCE identifies combinations of perturbations applied on a GRN or signaling network to trigger a shift from an undesired to the desired cell fate (Figure 2a). The core of the pipeline is (1) the application of the FVS control to identify reprogramming targets and (2) an attractor landscape estimation coupled with machine learning methods to predict the precise perturbations that drive the system from an initial state (that would lead to an attractor associated with an undesired phenotype) and towards an attractor associated with the desired phenotype.

**Fig. 2.**
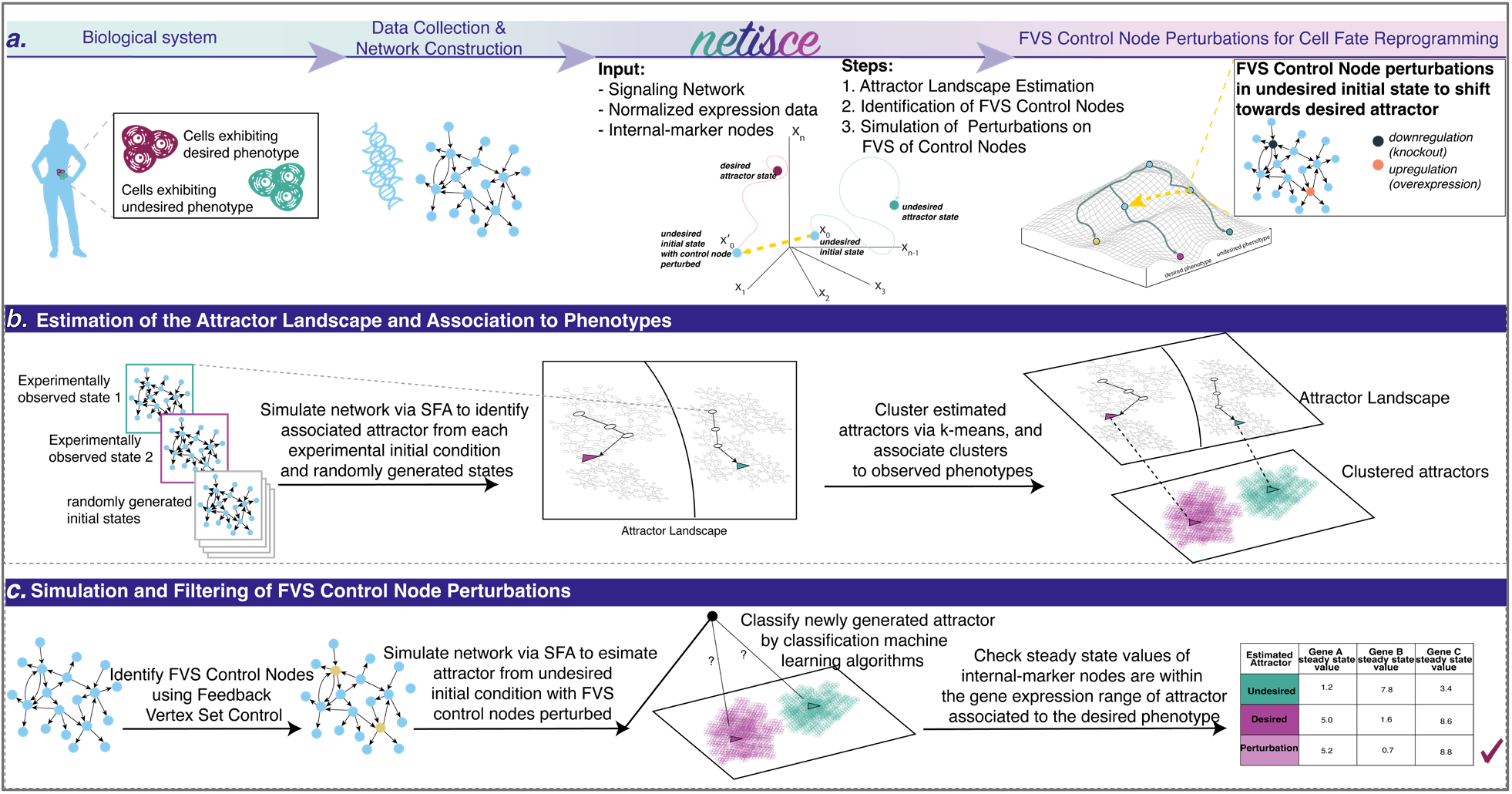
NETISCE Pipeline and Method Overview. **a** Researchers can collect data from cells exhibiting desired and undesired phenotypes and construct the signaling and regulatory networks governing cell reprogramming processes. The outputs of NETISCE are the combinations of network perturbations that shift the system from an undesired to a desired phenotype. **b** With the Signal Flow Analysis (SFA) algorithm^47^, in the first step, attractors are estimated by simulating the network with the initial states from normalized expression data and randomly generated initial states. The attractors are clustered via k-means, and the clusters are associated with desired (purple) and undesired (green) phenotypes. **c** In the second step, FVS control nodes are identified using an FVS-finding algorithm^55^. Perturbations on FVS control nodes are performed by setting the initial states of the system to the gene expression value of the undesired phenotype and overriding the states of FVS control nodes. In the third step, the sets of perturbations on FVS control nodes that achieve the desired reprogramming are identified using two filtering criteria. In the first criterion, the attractors generated from the perturbations on FVS control nodes are filtered using machine learning classification algorithms to obtain a set of perturbations whose attractors shifted from the cluster associated with the undesired phenotype to the cluster associated with the desired phenotype. In the second criterion, for user-defined internal-marker nodes, the steady-state expression values of the perturbations that passed the first filtering criterion (light purple) are evaluated to determine if their values have shifted to the gene expression range of the attractors associated with the desired phenotype.

### Method’s Validation

We provide here three application examples of NETISCE in experimentally validated studies of cell fate reprogramming. In addition, we present a comparative study of NETISCE applied to the *Drosophila melanogaster* patterning specification system described *in silico* both as Boolean and Ordinary Differential Equations Models (Supplementary Text 2, Supplementary Tables 1,2, Supplementary Figure 2).

### Reproducing experimentally validated perturbations to FVS control nodes for Cell Fate Specification in Ascidian Embryos

Using a GRN of cell fate specification in ascidian embryos, Kobayashi and colleagues experimentally verified that concerted perturbations to the network’s FVS could induce embryonic cells to the epithelial, mesenchymal, endodermal, notochord, brain, and pan-neural, and muscle tissue fates^52,53^. We performed simulations of the experimentally verified perturbations on FVS control nodes for cell fate specification in ascidian embryos using SFA.

The ascidian embryo GRN contained 92 nodes and 329 edges (Figure 3a). We identified all 26 FVSes within the ascidian embryo GRN, including the set of six FVS control nodes experimentally tested by Kobayashi *et al*.^53^: Foxa.A, Foxd, Erk Signaling, Neurog, Tbx6-r.b, and Zic-r.b. Without available normalized expression data, we simulated *in silico* unperturbed embryonic development by setting the initial activities of two genes necessary for normal embryonic development, Gata.a and Zic-r.a^54^, to 1 (the activated state) and all other nodes to 0 (inactivated state). The attractors of the unperturbed state and the seven experimentally validated combinations of perturbations (synchronous overexpression and knockout simulations) on FVS control nodes that induced the embryonic tissue fates were estimated using SFA. To evaluate the perturbations, we analyzed the steady-state expression values of the seven internal-marker nodes (one marker for each tissue) representing genes measured in the experimental study (Figure 3b, Supplementary Table 3). Attractors estimated by SFA can be compared analogously to the logarithm of the fold-change (*log*_2_*FC*) in differential gene expression analysis^47^, where the difference between the steady-state expression value of a node in attractors generated from different initial states or perturbations indicates that the gene expression of the node is upregulated if the difference is positive, or downregulated if the difference is negative (see Methods for details).

**Fig. 3.**
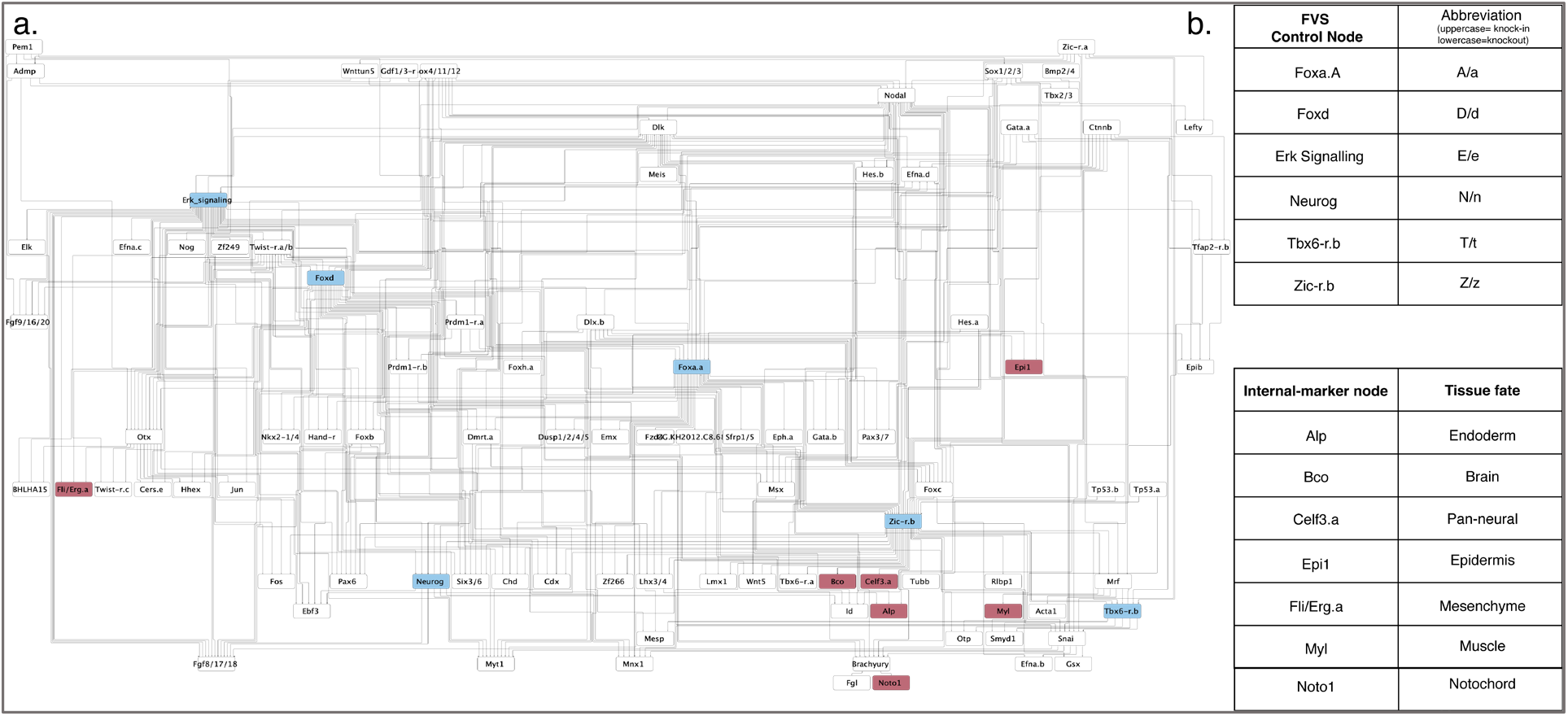
Cell Fate Specification in Ascidian Embryo Model. **a** Gene regulatory network of cell fate specification in the Ascidian Embryo *ciona intestinalis* from Kobayashi *et a.l*^53^. The network contains 92 nodes and 329 edges. Nodes highlighted in blue are FVS control nodes. Nodes colored in magenta are the internal-marker nodes used to evaluate if the perturbations on FVS control nodes successfully specified the desired cell fate. **b** The six control nodes identified by Kobayashi *et al.* that comprise the network’s minimal Feedback Vertex Set (FVS). Uppercase or lowercase abbreviations for each node indicate up or downregulation of the FVS control node in a given combination of perturbations (see Figure 4). These nodes are used as the FVS control nodes in our simulations. **c** The seven internal-marker nodes and the respective tissue fates where they are upregulated as identified by Kobayashi *et al.* experimental studies. These nodes were also used to identify successful perturbations in our simulations.

We successfully reprogrammed the unperturbed embryo to six of the seven tissue fates using the corresponding experimentally verified perturbations on FVS control nodes, giving the SFA in our pipeline an overall 85% accuracy (Figure 4, Supplementary Table 4). We could not reproduce reprogramming to the pan-neural tissue fate. However, the internal-marker node for the pan-neural cell fate (Celf3.a) was upregulated when we simulated the perturbation on FVS control nodes that experimentally induced the brain+pan-neural tissue fates.

**Fig. 4.**
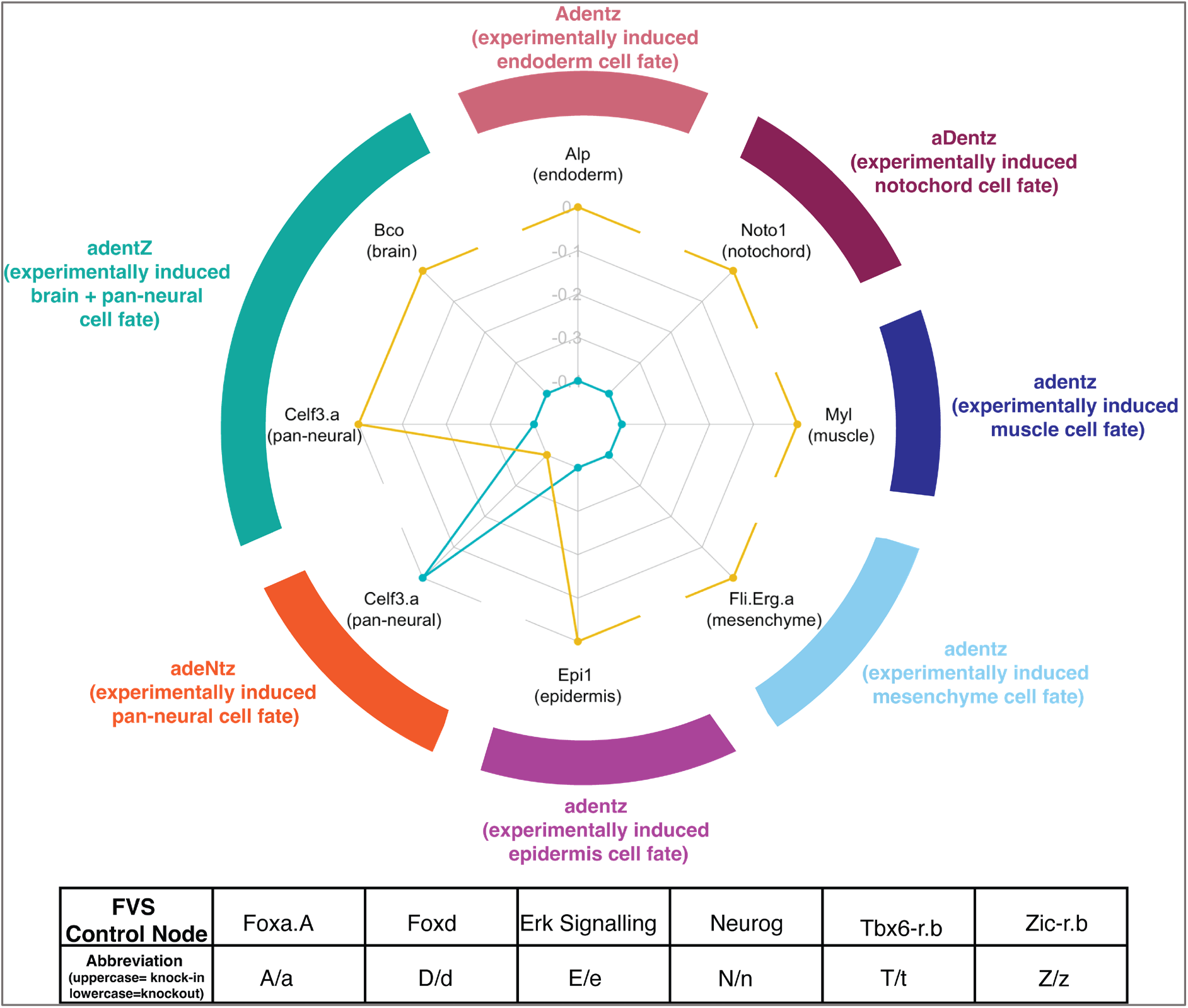
SFA simulations of perturbations on FVS control nodes for cell fate specification in Ascidian Embryos. Kobayashi *et al*.^53^ performed experimental combinations of perturbations on FVS control nodes to induce seven tissue fates in ascidian embryos. We aimed to reproduce these results *in silico* by simulating the combinations of perturbations on FVS control nodes using SFA. The results of each combination of perturbations are displayed in the radar plot. Each axis on the radar plot displays the steady-state expression value for an internal-marker node, with the tissue each internal-marker node represents in parentheses. The teal polygon shows the steady-state expression values of the internal-marker nodes in the attractor associated with the unperturbed state. The yellow polygon displays the internal-marker node steady-state expression values produced by applying one combination of perturbations on FVS control nodes to the unperturbed initial state. Note that the precise steady-state values have been scaled for plotting, and the raw data can be found in Supplementary Table 3. The outer colored ring denotes the combination of perturbations on FVS control nodes performed with the respective results displayed at that axis. The tissue fate that was induced experimentally by a perturbation is denoted in parentheses. Each letter and capitalization stands for a separate FVS control node and its perturbed state, as described in the table at the bottom. For a simulation of the perturbations on FVS control nodes to be considered successful, the steady-state expression value of the internal-marker node must be greater in the attractor produced by the perturbation on the FVS control nodes than the steady-state expression value in the attractor associated with the unperturbed state (the yellow polygon extends out past the teal polygon on the radar plot). We reproduced the cell fate specification results for 6/7 cell fates, excluding adentZ inducing the pan-neural cell fate.

### Identification of perturbations on FVS control nodes for induced pluripotent stem cell reprogramming from primed to naïve pluripotency

The mechanisms that maintain stem cells’ pluripotency or signal development are complex. Yachie-Kinoshita and colleagues constructed a Boolean Model to study pluripotent cell fate transitions under the constraint of signaling inputs^56^. The system was simulated under a stochastic asynchronous updating scheme. They simulated perturbations to every network node and identified targets to reverse primed pluripotency to naïve pluripotency. Yachie-Kinoshita *et al*. experimentally validated that the predicted targets reprogrammed primed pluripotency epiblast stem cells (EpiSCs) towards a naïve pluripotency embryonic stem cell (ESC) state. We used NETISCE to identify combinations of perturbations on FVS control nodes that reprogram EpiSCs towards the ESC state (Figure 5a).

**Fig 5.**
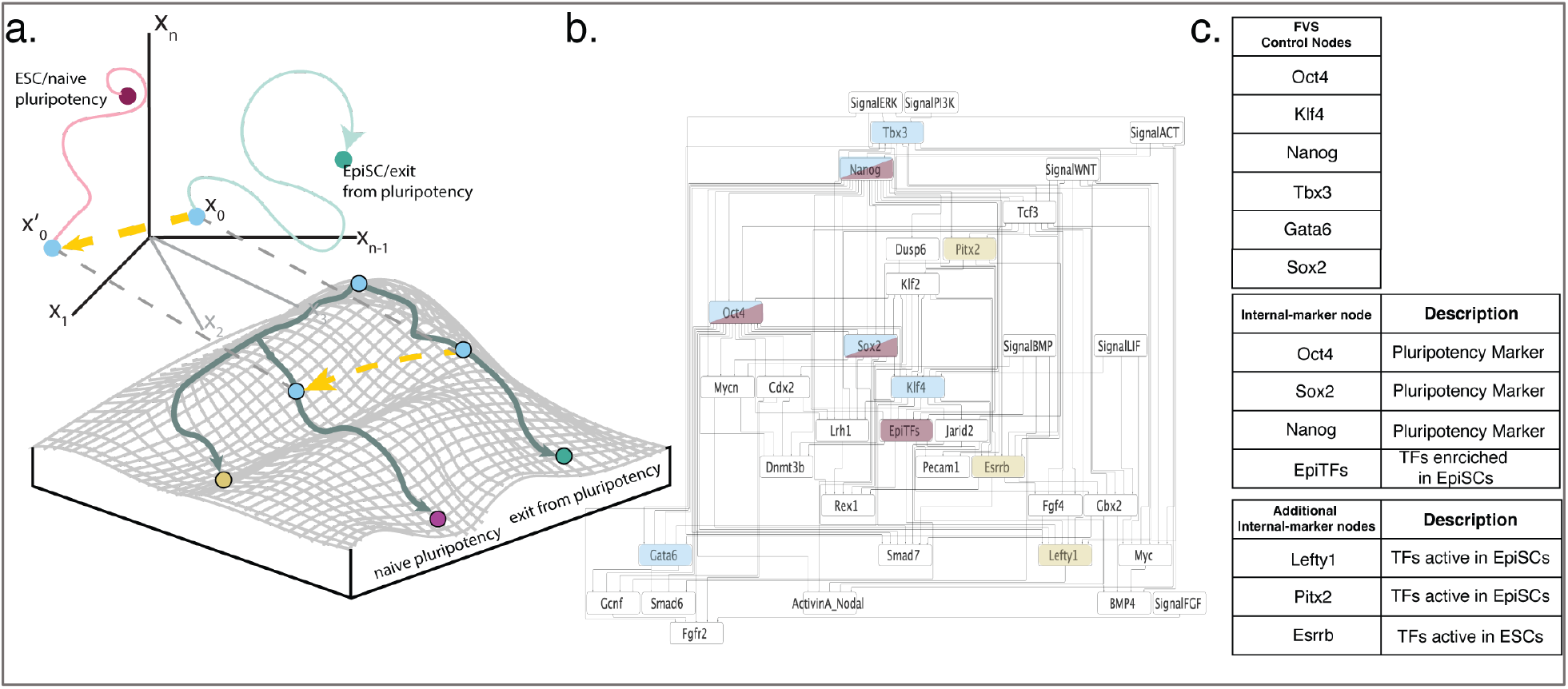
Reprogramming of Epiblast stem cells (EpiSCs) to Embryonic Stem Cells (ESCs). **a** The goal of this study is for NETISCE to identify the perturbations on FVS control nodes that shift EpiSCs towards ESCs, reverting from a primed to a naïve pluripotent state. To that end, we initialize the system with the gene expression data from EpiSC cells (x_0_), and, with the correct combination of perturbations on FVS control nodes, shift the system to a state (yellow arrow leading to *x*’ _0_) that leads to the ESC cell fate (magenta circle). **b** Pluripotent Stem Cell signaling network as constructed by Yachie-Kinoshita *et al*.^56^. The network contains 36 nodes and 143 edges. Nodes colored in blue are FVS control nodes, nodes with magenta coloring are the internal-marker nodes used by Yachie-Kinoshita *et al*., and gold nodes are the additional internal-marker nodes we used for further selection of perturbations on FVS control nodes. Note that the nodes Sox2, Nanog, and Oct4 are FVS control nodes and internal-marker nodes, and thus colored with both blue and magenta. **c** Key nodes for the desired cell fate reprogramming. There are six FVS control nodes within the system. Four internal-marker nodes were used by Yachie-Kinoshita *et al*. to identify successful reprogramming targets for the ESC state. We identified an additional three internal-marker nodes to filter perturbations on FVS control nodes. In the second column for the internal-marker nodes, we denote which behavior they represent in pluripotency signaling.

The pluripotency signaling network contained 36 nodes and 143 edges (Figure 5b). Using NETISCE, we estimated the six attractors from EpiSC and ESC gene expression data and attractors from 100,000 randomly generated initial states for a total of 100,006 attractors. On these 100,006 attractors, we performed k-means clustering. The optimal number of clusters identified by the elbow and silhouette metrics was k=2. One cluster contained the attractors generated from the initial state values of the EpiSC cells’ gene expression. The second cluster included the attractors generated from the ESC cell gene expression initialization.

NETISCE identified only one FVS in the network, comprising six nodes: Nanog, Oct4, Klf4, Sox2, Gata6, and Tbx3 (Figure 5c). Then, we simulated the 729 combinations of perturbations on the FVS control nodes. Of the 729 perturbations, 375 passed the machine learning classification filtering criterion. We identified each machine learning classification algorithm’s top 10 percent ranked features through feature importance analysis. Two out of the three top SVM features were FVS nodes. The top features across all three classification algorithms were mutually exclusive, alluding to their complementary nature (Supplementary Text 3, Supplementary Table 6). We used the same four internal-marker nodes as Yachie-Kinoshita *et al*.: Oct4, Sox2, and Nanog as markers of naïve pluripotency, and EpiTFs as the marker of enriched transcription factors in EpiSCs. The steady-state expression values of the internal-marker nodes in 132 of the 375 attractors calculated from the perturbations on FVS control nodes were in the range of gene expression values of the ESC associated attractors and thus passed criterion 2 for all replicates. Notably, one perturbation on FVS control nodes that passed both filtering criteria — overexpression of Nanog — was also identified and experimentally validated by Yachie-Kinoshita *et al*. (Figure 6a). In the Boolean simulations and experimental validation by Yachie-Kinoshita *et al*., Klf4 overexpression induced the ESC fate. Although Klf4 was an FVS control node in the network and its overexpression passed the machine learning filtering criterion, the perturbation did not pass the internal-marker node filtering criterion (Figure 6a). In this case, the steady-state expression values of Nanog, Sox2, and Oct4 (when considered as internal-marker nodes) did not reach the gene expression levels of the ESC state (Figure 6a). Overall, we show that overexpression of the FVS control node Nanog results in cell fate reprogramming to naïve pluripotency, in agreement with the results from the Boolean Model simulations and experimental validations.

**Fig. 6.**
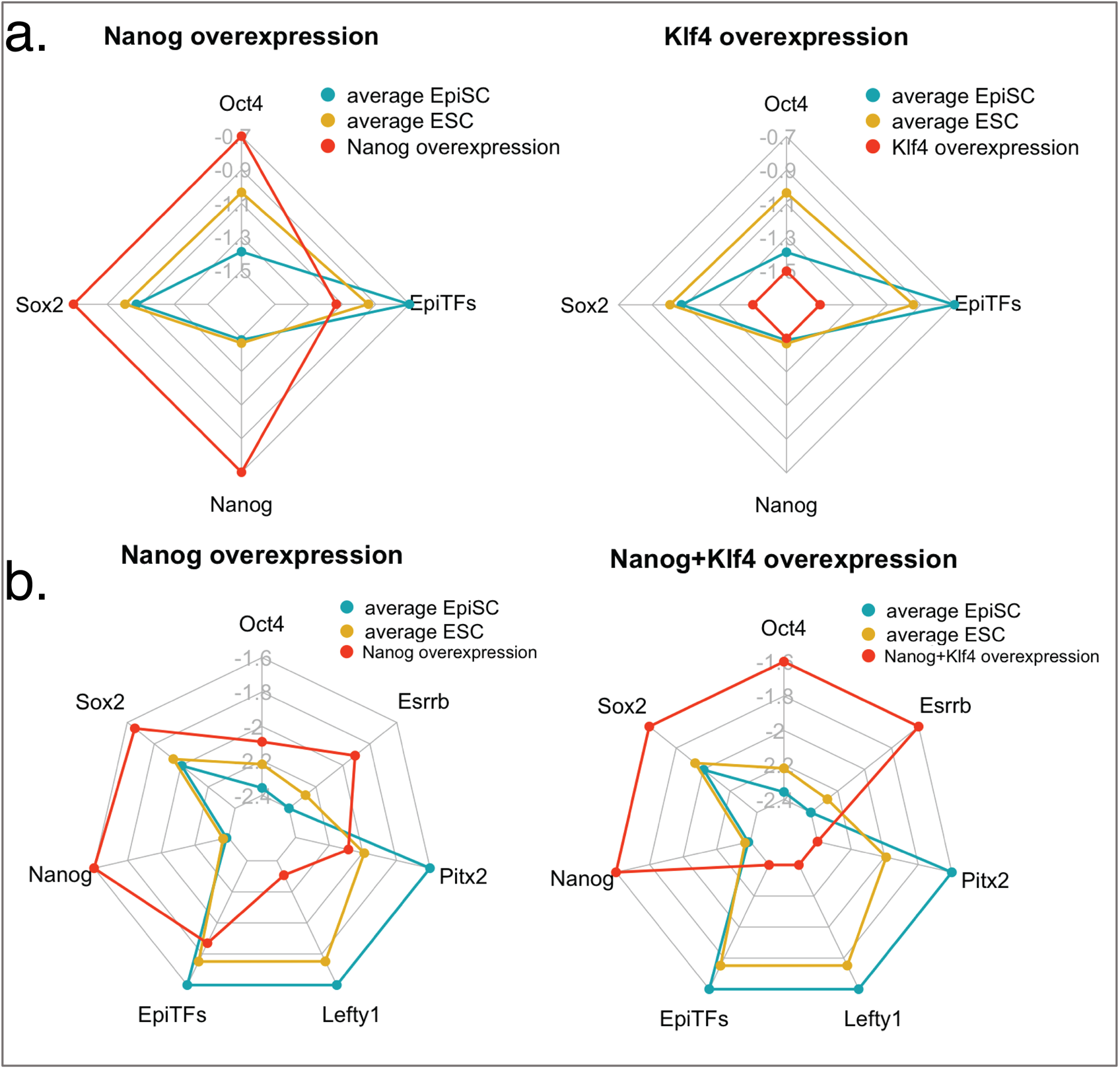
Results of simulations of perturbations on FVS control nodes for EpiSC reprogramming to ESCs. Radar plots for SFA simulations of perturbations on FVS control nodes in the pluripotent stem cell model. The title of each plot indicates the perturbation on FVS control nodes. Each axis on the radar plot displays the steady-state expression value for an internal-marker node. The blue polygons are the average steady-state expression values of the internal-marker nodes for attractors generated from the EpiSC experimental samples. The yellow polygons are the average steady-state expression values of the internal-marker nodes for attractors generated from the ESC experimental samples. The orange polygons are the steady-state expression values of the internal-marker nodes in the attractor produced by the specified perturbation on FVS control nodes. A perturbation is considered successful if 90% of the internal-marker nodes have steady-state expression values within the gene expression range of the ESC-associated attractor. In other words, if the yellow polygon at the axis of a specific internal-marker node extends beyond the blue polygon, the orange polygon must extend beyond the yellow polygon at that internal-marker axis. Conversely, if the yellow polygon at the axis of a specific internal-marker node does not extend past the blue polygon, the orange polygon must not extend past the yellow polygon at that internal-marker axis. **a** Radar plots using the four internal-marker nodes used in Yachie-Kinoshita *et al*. for the perturbations of Nanog overexpression and Klf4 overexpression. Klf4 overexpression does not pass the internal-marker node filtering criteria because the steady-state expression values of Oct4, Sox2, and Nanog are not within the range of expression of these genes in the attractor produced from ESC gene expression data. **b** Radar plots using the four internal-marker nodes used in Yachie-Kinoshita *et al*., and the additional three internal-marker nodes identified from differential gene expression data for two successful perturbations — Nanog overexpression and combined Nanog+Klf4 overexpression.

We explored the ability to further filter the perturbations on FVS control nodes by increasing the number of internal-marker nodes. We identified three additional nodes from gene expression data provided by Yachie-Kinoshita *et al*. These included Lefty1, Pitx2 (transcription factors active in EpiSCs), and Esrrb (a transcription factor active in ESCs). This reduced the 132 perturbations that passed filtering criterion 2 to 15 perturbations. Nanog upregulation was present in all 15 perturbations (Supplementary Figure 3). Nanog overexpression and the combination of Nanog+Klf4 overexpression were two of the fifteen perturbations that passed the internal-marker node criteria (Figure 6b, Supplementary Table 5). The combination of Nanog+Klf4 was not previously identified by Yachie-Kinoshita *et al*. However, the combination Klf4 and Nanog overexpression may play an essential role in maintaining pluripotency, as Klf4 links extracellular signaling information to positively regulate downstream Nanog transcription, and overexpression of Nanog was found to rescue pluripotency in the case of Klf4 knockdown^57^. Finally, based on the results of NETISCE, we identified a potential error in the underlying network and performed revisions to the network structure to improve simulation results (Supplementary Text 4).

### Identification of perturbations on FVS control nodes to overcome adaptive resistance to targeted MAPK inhibitor therapy in colorectal cancers

BRAF inhibition (BRAFi) therapy is a form of MAPK inhibitor (MAPKi) therapy used to treat cancer patients with mutant BRAF. BRAFi inhibits the MAPK signaling pathway, suppressing proliferation and inducing apoptosis. In colorectal cancers (CRCs), adaptive resistance emerges against BRAFi through the activation of MAPK signaling by upstream regulator EGFR. Park *et al*. were interested in identifying a gene that, when perturbed in combination with BRAFi, could prevent the development of adaptive resistance to BRAFi^58^. However, instead of inhibiting upstream molecules like in the BRAFi + EGRFi treatment, they searched for a target within the MAPK signaling pathway that could sensitize HT29 CRC cells to MAPKi therapy. By constructing a Boolean model of signaling pathways in CRC and simulating perturbations to every node in the model under an asynchronous updating scheme, Park *et al*. showed that BRAFi combined with SRC inhibition (SRCi) prevented the development of adaptive resistance via inhibition of ERK (MAPK1), a member of the MAPK signaling pathway. This result was validated experimentally in HT29 CRC cells. Therefore, with only the CRC signal pathway network structure, RNA-seq data from untreated HT29 cells, and functionally annotated mutational information as input to NETISCE, we sought to identify perturbations on FVS control nodes in combination with BRAFi that overcome adaptive resistance to MAPKi therapy (Figure 7a).

**Fig. 7.**
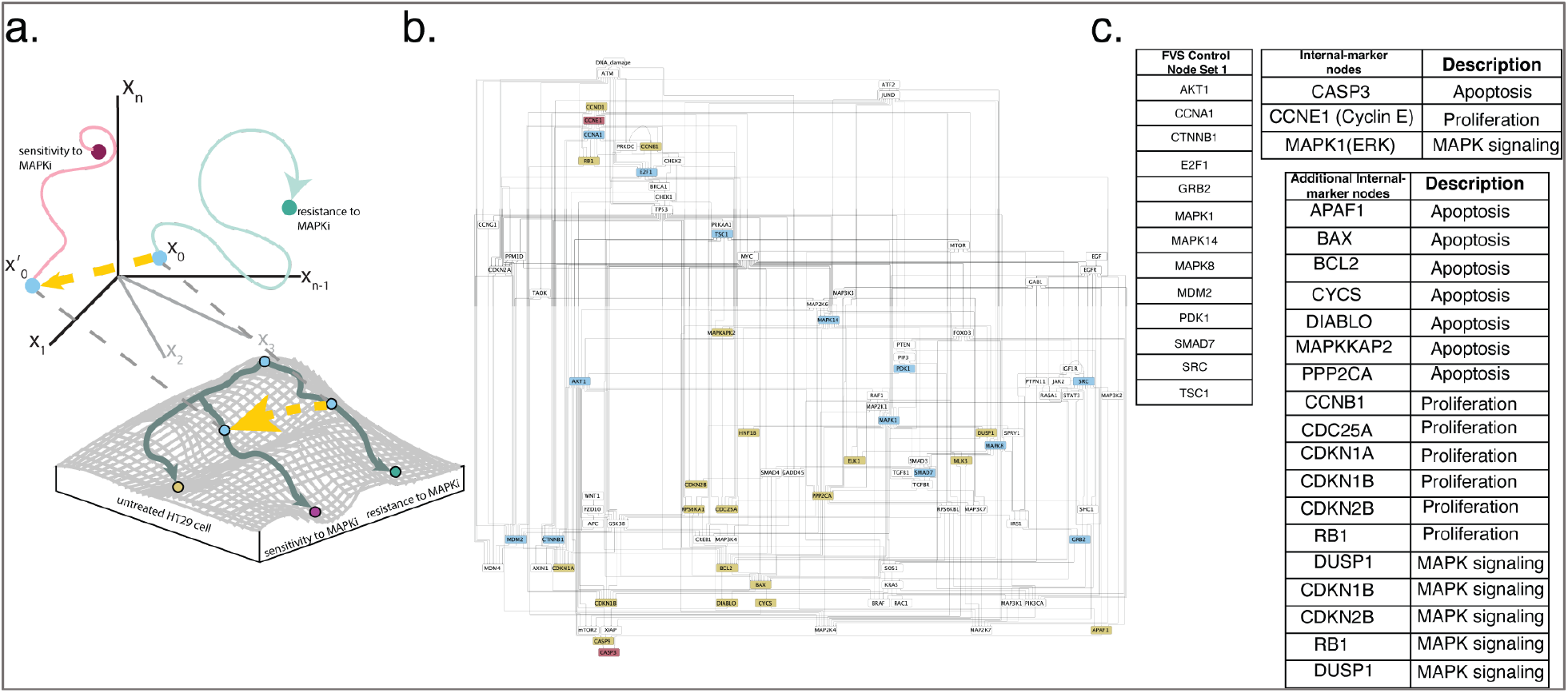
Overcoming adaptive resistance in the Colorectal Cancer model. **a** In this study, we use NETISCE to identify perturbations on FVS control nodes that shift BRAFi-treated HT29 cells away from a state of resistance to MAPK inhibitor (MAPKi) therapy and towards a MAPKi therapy sensitivity. We initialize the system with gene expression data from untreated HT29 cells (*x*_0_) and simulate BRAFi together with perturbations on FVS control nodes to shift the system to a state (yellow arrow leading to *x*’ _0_) that leads to sensitivity to MAPKi therapy (magenta circle). **b** Colorectal cancer signaling network introduced in Park *et al*.^58^. Blue nodes are control nodes, magenta nodes are internal-marker nodes, and gold nodes are additional internal-marker nodes used for further filtering of perturbations of FVS control nodes. **c** Key nodes for the cell fate reprogramming. One of the 68 FVS control node sets within the network is presented here, called *Set 1*. There are three internal-marker nodes used by *Park et al*. to identify targets to reprogram the system towards the MAPKi therapy-sensitive state. The three nodes are used to measure the phenotypes of apoptosis, proliferation, and MAPK signaling activity within the perturbed system. We identified an additional 17 internal-marker nodes to filter perturbations on FVS control nodes. The phenotypes associated with the internal-marker nodes are denoted in the second column.

Park and colleagues built a CRC signaling network containing 95 nodes and 337 edges (Figure 7b). We adapted the PROFILE method^59^ to verify that the generic CRC network and our SFA simulations preserved the phenotypic signatures of apoptosis and proliferation (see Supplementary Text 5 and Supplementary Figure 4).

The normalized gene expression data from an untreated HT29 sample was used for the initial activities for all SFA simulations, as data for HT29 cells with BRAFi and BRAFi+EGFRi treatment is unavailable. In HT29 cells, PIK3CA and BRAF have gain-of-function mutations, while APC, SMAD4, and TP53 have loss-of-function mutations. Therefore, states of nodes with gain-of-function/loss-of-function mutations were overridden to the appropriate overexpression or knockout state using our modified SFA equation for perturbations (see Methods). Next, we simulated the treatment of an untreated HT29 cell with BRAFi (HT29_BRAFi) or BRAFi+EGFRi (HT29_BRAFi+EGFRi) to obtain attractors related to the MAPKi therapy-resistant state and MAPKi therapy-sensitive state, respectively. These simulations used the normalized expression data of the untreated HT29 sample as initial state values and included the appropriate mutational overrides. To simulate BRAFi and EGFRi, the state of these nodes was overridden to a knockout state using the modified SFA equation for perturbations. The optimal k-means clustering of the attractors obtained from the untreated HT29, HT29_BRAFi, HT29_BRAFi+EGFRi treated initial conditions, and the attractors from 100,000 randomly generated initial states was k=3; as desired, the untreated-state, MAPKi therapy-resistant state and MAPKi therapy-sensitive state associated attractors were found in separate clusters. NETISCE exhaustively identified 68 FVSes in the CRC static signaling network; the union of all FVSes contained 25 nodes, and each FVS had a combination of 14 out of the 25 nodes. All FVSes contained SRC. In addition, each FVS included TP53, a loss-of-function mutant gene in HT29 cells whose state was already overridden to a knockout state in our simulations; therefore, additional perturbations to TP53 were not performed, reducing the number of FVS control nodes that could be perturbed to 13 (Supplementary Data 1).

We present the results of using one FVS, referred to as Set *1* (Figure 7c), to identify combinations of perturbations that shift the system from the MAPKi therapy-resistant phenotype to the MAPKi therapy-sensitive phenotype. We simulated 1,594,323 combinations of perturbations to the 13 FVS control nodes in *Set 1* to identify their corresponding attractors. First, 232,114 of the 1,594,323 attractors generated from the perturbations on FVS control nodes were classified to the MAPKi therapy-sensitivity associated cluster by at least two out of three machine learning classification algorithms. FVS control nodes or internal-marker nodes were part of the top 10 percent of ranked features revealed by the feature importance analysis on the three machine learning classification algorithms (Supplementary Text 6, Supplementary Table 7). Then, the 232,114 perturbations that produced the attractors that passed criterion 1 were filtered by the three internal-marker nodes that were used as output readout nodes in Park *et al*. (Figure 7c): CASP3 (apoptosis marker), Cyclin E (proliferation marker), and MAPK1/ERK (MAPK signaling activity marker). The internal-marker nodes in the attractors produced by 52,703 of the 232,114 perturbations on FVS control nodes had steady-state expression values in the range of gene expression values of the MAPKi therapy sensitivity associated attractors.

Notably, the combination of BRAFi + SRCi was the smallest perturbation set that passed both filtering criteria (Figure 8a).

**Fig. 8.**
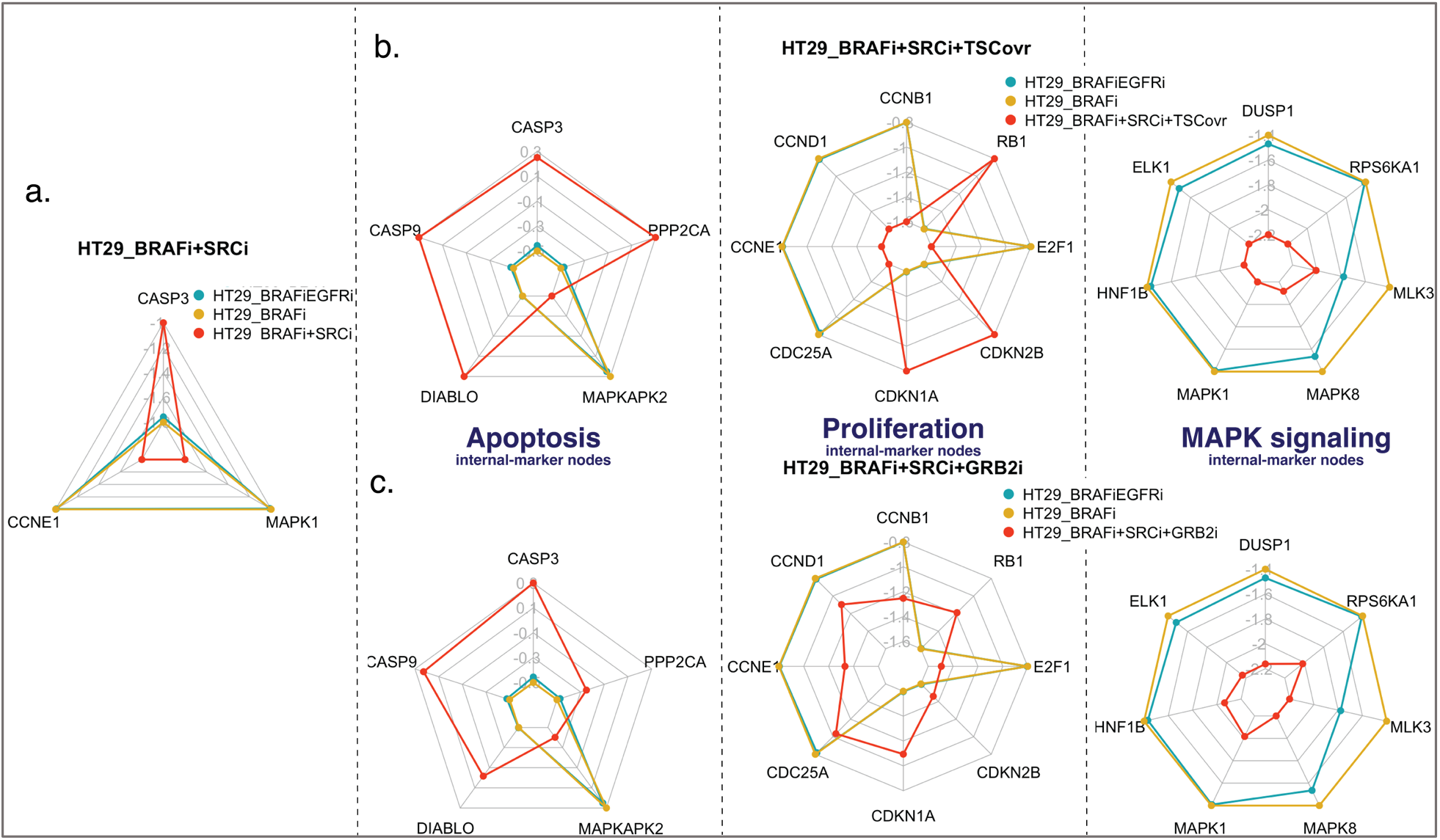
Results of simulations of perturbations on FVS control nodes for overcoming adaptive resistance in colorectal cancer. Radar plots for perturbations on FVS control nodes in HT29 cells in the colorectal cancer signaling network. The title of each plot indicates the perturbation on the FVS control nodes with BRAFi. Each axis on the radar plot displays the steady-state expression value for an internal-marker node. The blue polygons are the steady-state expression values of the internal-marker nodes in the HT29_BRAFi+EGFRi (MAPKi therapy-sensitive) associated attractor. The yellow polygons are the average steady-state expression values of the internal-marker nodes in the HT29_BRAFi (MAPKi therapy-resistant) associated attractor. The orange polygons are the steady-state expression values of the internal-marker nodes in the attractor produced by the specified perturbation on FVS control nodes with BRAFi. A perturbation is considered successful if 90% of the internal-marker nodes’ steady-state expression values are within the gene expression range of the HT29_BRAFi+EGFRi-associated attractor. **a** Radar plot of the three internal-marker nodes used by Park *et al*. for the simulated BRAFi+SRCi perturbation. The radar plot shows that the perturbation of BRAFi+SRCi increases apoptosis and decreases proliferation and MAPK activity based on the internal-marker nodes. **b** Radar plots for the two smallest sets of perturbations on FVS control nodes (SRCi+TSC1 overexpression and SRCi+GRB2i) with 20 internal-marker nodes, separated by the apoptosis, MAPK signaling, and proliferation phenotypes. The plots show that the internal-marker node values are within the ranges of the MAPK sensitivity associated phenotype for all internal-marker nodes.

We considered an additional 17 internal-marker nodes related to apoptosis, proliferation, and MAPK signaling (Figure 7c). Of the 52,703 perturbations on FVS control nodes, 1,266 met the filtering threshold of 90% for the apoptosis, proliferation, and MAPK internal-marker node steady-state values. SCRi was present in all combinations of control nodes that passed the two criteria (Supplementary Figure 5).

The smallest sets of perturbations were two pairs of two control nodes. The first reprogramming pair consisted of SRCi and TSC1 overexpression (SRCi+TSC1ovr) (Figure 8b), while the second reprogramming pair was comprised of SRCi and GRB2i (SRCi+GRB2i) (Figure 8c). TSC1 overexpression has been studied in the context of adaptive resistance to MAPKi. TSC1 promotes cell death by inhibiting mTOR activity, and mTOR inhibition in combination with BRAFi has been shown to overcome adaptive resistance in BRAF-mutant melanoma^60^. However, a complex of TSC1 and TSC2 can be inactivated by ERK phosphorylation, leading to increased mTOR signaling^61^. In the context of CRC, GRB2 is an important protein for transmitting oncogenic signaling and promoting tumorigenesis and metastasis^62^. Interestingly, the protein Gab2, a binding partner of GRB2, was directly upregulated by BRAFi in BRAF mutated cancers^63^. Impairing the interaction between GRB2 and Gab2 sensitized cells to BRAFi therapy and prevented additional oncogenic signaling and metastasis in HT29 cells^63,64^.

### Evaluating the Robustness to Noise of NETISCE

Commonly, gene expression data can be infiltrated with extrinsic noise from experimental methods or biological variability. We evaluated the robustness to noise of NETISCE to predict perturbations on FVS control nodes with noisy experimental gene expression data. Our approach uses COPASI^65^ to simulate two differential equation models of cell reprogramming and add different noise levels into some initial states. We use these generated initial states to simulate 1,000 triplicates of the desired and undesired phenotype normalized gene expression data per noise level varying from 0% to 50% (see Methods). Overall, applying this analysis in the ODE model of Drosophila Segment Polarity genes^66^ and a stochastic DE (SDE) model of cell fate differentiation in pancreatic cells^35^, we found that NETISCE is strongly robust to up to 50% of noise for which more than 75% of the perturbations on FVS control nodes were still correctly identified.

The ODE model of the Drosophila segment polarity GRN was used in FVS control studies. The authors identified a perturbation on the FVS that shifted the system from the unpatterned to the wild-type trajectory (Supplementary Text 2). In this model, NETISCE correctly predicts the specified perturbations on FVS control nodes that shift the system from the unpatterned to the wild-type phenotype in over 92% of the simulations for the 1,000 triplicates with 30% noise and 75% of simulations for the triplicates with 50% noise (Supplementary Table 8).

The Zhou *et al*. SDE model of pancreatic cell fate differentiation was used to simulate the reprogramming of pancreatic exocrine cells to beta-cells^35^. We identified 31 combinations of perturbations on FVS control nodes that could reprogram exocrine cells to beta-cells (Supplementary Figure 7, Supplementary Text 7). For more than 89% of the simulations of 1,000 triplicates with up to 20% of noise, NETISCE correctly predicted the 31 combinations of perturbations on FVS control nodes (Supplementary Table 9). At 50% noise in the initial states, 84.1% of the simulations for the 1,000 triplicates correctly predict the 31 combinations of perturbations on FVS control nodes.

## DISCUSSION

With the rise in availability of multi-omics datasets and tools for constructing gene regulatory and intracellular signaling networks from these data, there is a growing need for cell reprogramming methods that are data-driven and amenable for larger-scale biological networks where parameters to model all the system components may not be available or be challenging to estimate. We have developed NETISCE, a tool that identifies cell fate reprogramming targets using the FVS control, attractor landscape estimation, and machine learning methods. By reproducing experimental and mathematical model results, we show that NETISCE can identify cell fate reprogramming targets and their perturbations using the system’s static network, gene expression data from the undesired cell phenotype, and a set of nodes used as internal-marker nodes for the desired and undesired phenotypes.

NETISCE offers a unique approach to identifying cell fate reprogramming targets through its application of control theory and a dynamical systems-based framework. First, by employing the structure-based FVS control, we consider feedback loops that commonly regulate biological functions essential for identifying reprogramming targets. Since our target search focuses on FVS control nodes, we guarantee that the identified nodes are sufficient for cell fate reprogramming. Applying the FVS control to the problem of identifying cell fate reprogramming targets has distinct advantages. In comparison to network-based approaches, we do not need to compare the structure of multiple networks to identify cell fate reprogramming targets; in contrast to dynamical systems-based methods that do not apply control theory, full dynamical information for all network components is not required, nor is there a need to screen all network elements.

The FVS control contrasts with other control theory approaches for identifying targets like Data Guided Control, which may fail to capture cell reprogramming dynamics due to its linear assumption of regulatory dynamics^41^. It is important to note that FVS control for identifying reprogramming targets requires a high-fidelity static network. As Mochizuki and colleagues observed^5^ and later, Kobayashi and colleagues showed experimentally^52,53^, if a cell state cannot be reached by perturbations to the FVS control nodes, then the FVS was not correctly identified, and network revision should be performed.

Secondly, the SFA algorithm and machine learning methods allow us to identify the specific perturbations on FVS control nodes required for cell fate reprogramming by estimating system dynamics and the attractor landscape. In our approach, by associating phenotypes to the attractor landscape via k-means clustering and classifying the attractors produced by perturbations on FVS control nodes via machine learning classification methods, we observe when the system has shifted towards the attractor associated with the desired phenotype. Lastly, we have modified the original SFA algorithm^47^ to apply FVS control-based perturbations. Unlike the SFA control method introduced by Lee and Cho^47^, where edge modifications (removals or additions) were implemented to perform perturbations, our method of perturbations to a node’s state maintains the FVS of the network. Additionally, we have implemented permanent overrides on FVS control nodes rather than the original form of SFA perturbations that are transiently applied by changing only the initial states of nodes. Transient perturbations are not applicable in the context of FVS control. First, as defined by Fielder *et al*. and Mochizuki *et al*., overrides to the states of the FVS control nodes guarantee that the system will arrive at any desired attractor^5,6^. Second, our implementation mimics the experimental perturbations in Kobayashi *et al*., where FVS genes were permanently overexpressed or knocked out in the ascidian embryo. While transient perturbations could be used to simulate some types of single-drug treatments, their use in the context of the FVS control may not produce the desired reprogramming. In mathematical models, perturbations on FVS control nodes can drive the system to fixed-point attractors or cycles. However, the SFA algorithm currently identifies only fixed-point attractors due to its tolerance threshold. A future update could modify the tolerance threshold to identify limit cycles.

NETISCE reproduced the simulated and experimentally validated results in different applications of cell fate reprogramming. We have shown that NETISCE reproduced the results of *in silico* simulations of the FVS control in both Boolean and ODE models and, that it is highly robust to noise. Finally, although NETISCE performs synchronous, deterministic simulations, we reproduced results from asynchronous and stochastic mathematical models.

Importantly, we have shown that NETISCE can be used to personalize simulations on a network when provided with expression (and, if available, mutational profiles) data for a specified sample. We use PROFILE^59^ to verify the network structure and SFA simulations using patient tumor data on the CRC signaling network. PROFILE can be applied in analogous cancer studies and when the signatures of interest are specific genes related to a biological function or phenotype, such as those in Molecular Signatures Database^67^. However, there is not an established model verification method for problems outside of cancer. In these problems, such as the iPSC example, NETISCE simulations and feature importance analysis can evaluate the correctness of the network and potentially correct errors if literature information or data exists (see Supplementary Text 4).

In the example of cell fate specification in ascidian embryos, our SFA simulations succeeded in reproducing 85% of the experimentally validated perturbations. This accuracy may be a limitation of the SFA estimation algorithm. The biological process of pan-neural tissue specification may explain the inability to induce the pan-neural tissue fate by Neurog overexpression. Otx – a gene downstream of Neurog involved in pan-neural cell fate specification – is inactive in the ascidian embryo until the 32-cell stage of development^54^. Since overexpression of Neurog was performed at the start of the simulation, we may not be accurately simulating the timing of its pan-neural inducing effect. A future update to the SFA algorithm could perform asynchronous stochastic simulations, allowing for time-delayed perturbations and potentially producing information regarding the specific timings of perturbations for successful cellular reprogramming^41,68^.

Our FVS control-guided method reduces the number of simulations needed to be performed to identify targets for cell fate reprogramming. In the reprogramming tasks for the pluripotent stem cells and CRC problems, some of the perturbations to the FVS control nodes that NETISCE identified as successful were a subset of the perturbations found in Boolean models, where reprogramming targets were identified by simulating perturbations to every node in the system. In addition, NETISCE revealed combinatorial strategies for cell fate reprogramming in both models. In the case of the pluripotent stem cell model, Nanog is essential to maintain pluripotency, and additional overexpression of Klf4 could make the reprogrammed cell unreceptive to extracellular signaling that may signal to exit from pluripotency, preserving high levels of Nanog activity^57^. However, unlike in the Boolean model simulations, we could not identify that Klf4 overexpression alone could reprogram cells to the ESC cell fate. This result may be because SFA does not capture cooperative activity in the same manner as Boolean logic; as in SFA, the initial activity of a node can influence the state of a node at each time step. In the model of adaptive resistance to MAPKi therapy in CRC, the combination of BRAFi+SRCi+TSC1ovr could further increase sensitivity to treatment and the rate of apoptosis by MAPK and mTOR signaling inhibition^61^. Alternatively, BRAFi+SRCi+GRB2i can increase sensitivity to MAPKi therapy and prevent the metastatic spread of CRC tumors^63^.

With the potential to identify hundreds of thousands of perturbations that satisfy NETISCE’s filtering criteria, a method for prioritizing perturbations on FVS control nodes is essential. A simple method employed in NETISCE is generating a secondary set of internal-marker nodes to filter perturbations. The number of necessary internal-marker nodes is dependent on the user’s goals and the specific application. For example, a minimal set of marker nodes may generate thousands of successful perturbations on FVS control nodes, allowing the user to understand patterns of perturbations and potentially focus on specific network modules. However, a larger set of internal-marker nodes may reduce the number of successful perturbations on FVS control nodes, which could help identify predicted targets to be verified experimentally. Depending on the system, it may also be beneficial to prioritize the smallest combinations of perturbations on FVS control nodes to ease experiments or prevent off-target effects. Another prioritization approach could score perturbations based on the strength of their effect on the target phenotypes while minimizing side-effects, similar to the method implemented by Park *et al*. in the CRC Boolean model simulations^58^. A node received a high score if, when the probability of its activity in the results of the asynchronous stochastic simulations indicated inhibition, it would prevent adaptive resistance via ERK reactivation, promote the therapeutic side-effect of increased apoptosis, and prevent the adverse side-effect of increased proliferation. This scoring could be modified for NETISCE. Majorly, NETISCE would need to perform stochastic asynchronous simulations like the Boolean Model simulation framework used by Park and colleagues. The scoring also needs to consider overexpression, knockout, and combinations of perturbations on FVS control nodes. Next, the processing of internal-marker nodes could be modified to consider nodes related to side effects. Finally, an algorithm for path-finding and determining the effect of a perturbation on FVS control nodes on off-target nodes would need to be implemented.

In stem-cell reprogramming, where changes to the epigenetic profile are a significant factor in reprogramming efficiency^69^, implementing a scoring of combinations of perturbations on FVS control nodes that considers epigenetic information could be highly effective to rank reprogramming targets. This information, which is incorporated into the tool, IRENE for both GRN construction and scoring of potential transcription factor-reprogramming targets, increased reprogramming protocol efficiencies in some cases by more than 900%^37^. Similar to the method of Park *et al*., this would also require implementing a stochastic simulation framework and could only be considered if the epigenetic information was available for both the undesired and desired states.

Additionally, if applicable, such as in a disease model, information like druggability or drug synergies could be incorporated into prioritizing combinations of perturbations on FVS control nodes^70,71^. For example, PHAROS^71^, a meta-database of drug-target information, can be used to assess the druggability of SRC, GRB2, and TSC1. Currently, six drugs are approved SRC inhibitors. However, there are no approved drugs for GRB2 inhibition nor known drugs or molecules that bind to TSC1 and promote its overexpression. Therefore, SRCi would be a likely first candidate for preventing adaptive resistance in combination with BRAFi.

Our dynamical systems-based analysis using static biological networks and experimental data to estimate the attractor landscape and perform combinations of perturbations to control nodes provides a valuable tool for intracellular signaling analysis. Because NETISCE can be applied to biological networks of a larger scale that are not fully parameterized, we envision it as a primary tool for cell fate reprogramming studies. Experimentalists can use the results generated from NETISCE to prioritize wet-lab perturbation experiments. At the same time, mathematical modelers can focus model construction towards regions that appear to be more relevant to the desired reprogramming task and fine-tune reprogramming target predictions by exactly solving for attractors rather than estimating steady states. Finally, our method produces useful and potentially novel combinations of perturbations for cell fate reprogramming that could eventually be applied for treatments in disease models to recover healthy cell phenotypes in biological systems.

## METHODS

### NETISCE Overview

#### INPUT

There are three required inputs for NETISCE: (1) a static network representing a biological system, (2) a set of normalized gene expression data from cells with an undesired phenotype, and (3) a set of internal-marker nodes — user-defined nodes within the network that can be used as a point of reference to verify that their gene expression levels match the expected values in desired and undesired phenotypes. Normalized gene expression data for cells with the desired phenotype can also be provided (see the Pluripotent Stem Cell example) but is not required for all use cases, such as simulating adaptive resistance to treatment (see the Colorectal Cancer example). Optionally, the input can include mutational data to specify the rules for the network’s simulations.

#### Step 1. Estimation of the attractor landscape

The goal of the first step of NETISCE is to estimate the region of the attractor landscape containing steady-states associated with the desired and undesired phenotype (Figure 2b). The network is simulated using an adapted version of SFA^46^. SFA estimates signal flow, the information conveyed by a series of reactions as represented in a signaling network or GRN, based only on topological information in the network and an initial state of the network nodes. The output of SFA is the logarithm of the steady-state value, which we refer to as the steady-state expression value for each network node. The initial states of the network nodes are based on the normalized gene expression levels. We simulate the system using SFA for each provided experimental sample until reaching the attractor.

We generate randomly sampled initial states whose values fall in the ranges of the normalized expression values for each of the supplied phenotypes in the experimental data and apply SFA to compute a sufficiently large number of attractor states^34^. All the computed attractors are then clustered via k-means clustering. The elbow and silhouette metrics are calculated to determine an optimal k^51,52^. The clusters are also evaluated using the internal-marker node values.

#### Step 2. Virtual screenings on FVS control nodes

In this step, NETISCE identifies FVS control nodes and simulates combinations of perturbations to their activity (Figure 2c). First, the FVS control nodes are identified via a simulated annealing algorithm to determine the FVS of the network^53^. This step can be performed exhaustively to identify all FVSes in a network. Then, virtual screenings of combinations of perturbations on the FVS control nodes are performed using the SFA algorithm. In these simulations, the initial states of the network nodes are set to the normalized expression values of the cells with an undesired phenotype. We have modified the SFA pipeline to implement overrides to control nodes to simulate overexpression, knockout, or no change to a node’s activity.

#### Step 3. Filtering sets of perturbations on the FVS control nodes

The final step of the pipeline aims to identify the combinations of perturbations on the FVS control nodes that result in the desired cell fate reprogramming (Figure 2c). We employ two filtering criteria to evaluate the combinations of perturbations on FVS control nodes. The first criterion uses Random Forest, Support Vector Machine, and Naïve Bayes machine learning classification algorithms to classify the attractors generated by perturbations on FVS control nodes. In this classification step, using the previously clustered attractors by the k-means analysis, the attractors generated by perturbations on FVS control nodes are classified either in the cluster(s) associated with the undesired or desired phenotype^54–56^. To pass this filtering criterion, an attractor generated from the perturbation on FVS control nodes must be classified to the cluster associated with the desired phenotype by at least 2 out of 3 classification algorithms. After passing the first criterion, perturbations on FVS control nodes are evaluated by the second filtering criterion, which focuses on the steady-state expression values of the internal-marker nodes. Perturbations on FVS control nodes where at least 90% of the internal-marker node steady-state expression values are within the expression ranges of the attractors associated with the desired phenotype pass this filtering step. These criteria produce a final set of perturbations on FVS control nodes that are considered capable of reprogramming from an undesired cell fate towards the desired cell fate.

### Estimation of Steady-States using Signal Flow Analysis

Signal Flow Analysis (SFA) is based on the Signal Propagation algorithm developed by Lee and Cho^47^. The algorithm is a linear difference equation that computes the activity of a network node at a given time in terms of the state of the network node at the previous time step, the effect (activating or inhibiting), and the weight of the influence of its *m* incoming edges, and the initial state of the node. Precisely, the logarithm (*log*_2_) of the steady-state activity *x_i_* (*t* + 1) of a node *i* at time *t* + 1 is estimated by the initial state of the node and the activities of its regulators at time t using the following equation:

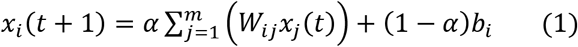

where *x_j_*(*t*) is the logarithm of the activity of *j*, a node connected to *i* by an incoming edge, at time *t*. The *W_ij_* is the weight of the edge between node *j* and node *i*, which represents how much influence node *j* exerts on node *i* through the edge. *W_ij_* is defined as:

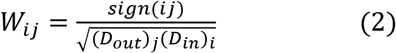

where *sign*(*ij*) is the value of the edge between *i* and *j* (1 for activating edges, −1 for inhibiting edges), *D_out_* is the out-degree of *j*, and *D_in_* is the in-degree of *i*. Finally, *b* is the logarithm of the initial state of node *i* and *α* is a hyperparameter used to weigh the influence of the network structure and initial node state on the Signal Flow. By default, in our pipeline, the hyperparameter *α* is set to 0.9 to provide greater weight to the network topology rather than the initial activity based on the parameter settings used in previous control studies using SFA^72^.

To identify an attractor of the system, the signal propagation equation is solved for all network nodes synchronously until the difference between *x*(*t* + 1) and *x*(*t*) is less than a tolerance threshold (by default, this tolerance threshold is 10^-6^).

The steady-state expression values of two attractors under different simulation inputs can be compared by computing the difference between two logarithms of the steady-state expression values produced from SFA, analogously to a logarithm of the fold-change (*log*_2_FC) in differential gene expression analysis^47^. Although the actual numerical value cannot be used to measure the magnitude of the change in expression, positive difference values indicate that the specified perturbation led to a shift in signal flow that increases the gene’s activity at the steady state. In contrast, negative values predict decreased activity at the steady state due to the perturbation. When estimating the attractor landscape, we begin by solving the signal propagation equation for the system using the experimental data for each sample; the initial activity of network nodes is set to the normalized expression values. In most cases, the number of experimental samples is insufficient for landscape estimation. Therefore, NETISCE can generate randomly sampled initial states (100,000 by default), whose initial state values are calculated from the ranges of the normalized expression values for each of the supplied phenotypes in the experimental data.

### Association of the Attractor Landscape Clusters to Experimental Phenotypes

We employ k-means clustering to partition the attractors estimated from the normalized expression data and the randomly generated initial states. We confirm that the attractors computed from the undesired and desired experimental samples are different. We use two metrics to determine the optimal number of k clusters. The first is the elbow metric, which determines the optimal k by finding the minimal intra-cluster variation^73^. The second is the silhouette metric, which aims to identify the optimal k as the number of clusters with minimal intra-cluster variation and maximal inter-cluster variations^74^. When the two metrics disagree on the optimal k, the smallest of the potential optimal k-values is chosen where the attractors estimated from the undesired and desired phenotype experimental samples do not appear within the same cluster(s).

Finally, we use the internal-marker nodes to confirm that the steady-state expression values of these nodes agree with experimental data or literature. NETISCE checks that the steady-state expression value of each internal-marker node in the attractors associated with the undesired phenotype and desired phenotype matches the expected differential gene expression patterns. In the scenario where only the experimental data for the undesired phenotype was provided for the initial states, NETISCE verifies that the cluster(s) containing the attractors generated from the experimental data only have attractors where the internal-marker node steady-state expression values are within the range of expression values of the undesired phenotype. If experimental data for both the undesired and desired phenotype is supplied for initial states, then NETISCE confirms that the attractors generated from the two phenotypes do not appear in the same cluster and that their internal-marker node steady-state expression values are within the appropriate expression value ranges. For example, consider a gene with a higher expression in cells in the desired phenotype than the undesired phenotype. NETISCE will verify that the steady-state expression value in the attractor associated with the desired phenotype is greater than the value in the attractor associated with the undesired phenotype (*i.e*., that the difference between the steady-state expression value for the internal-marker node in the attractor associated with the desired phenotype and the undesired phenotype is positive). Also, when multiple samples are given for each phenotype, NETISCE verifies that the steady-state expression values of the internal-marker nodes in the attractors of the undesired phenotype do not overlap with the values in the attractors of the desired phenotypes. If an overlap occurs, the internal-marker node is unreliable for analysis to separate the attractors in the different phenotypes. Thus, if the values of the internal-marker nodes do not match the literature or do not separate well between the attractors of the undesired and desired phenotype clusters, the user may elect to revise network structure, remove specific internal-marker nodes, or adjust simulation settings.

### Identification of the minimal Feedback Vertex Set

Structure-based methods study the controllability of systems based solely on the structure of the network^5,75,76^. In recent years, structure-based control methods for systems with non-linear dynamics have been proposed. One such structure-based control method for non-linear dynamics is the Feedback Vertex Set Control introduced by Mochizuki *et al*.^5,6^. Feedback Vertex Set Control is a structure-based control method focused on the controllability of the system by restricting the target states to attractors. Mochizuki *et al*. mathematically proved that for a network governed by non-linear dynamics like cell signaling, the control action of overriding the state variables of the feedback vertex set (FVS) into a targeted desired trajectory ensures that the system will asymptotically approach the desired trajectory. Consider a directed graph *G* = (*V,E*) comprised of node set *V* and edge set *E*. The node states of *G* are described by the ODE

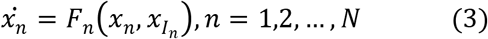

where for the dynamics *x* of node *n* ∈ *V, I_n_* is the set of nodes that regulate node *n,* such that self-regulatory loops (*n ∈ I_n_*) are only positive. Additionally, we assume *F_n_* satisfies decay condition:

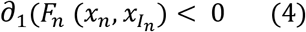

for all *n* where ∂_1_ is the partial derivative w.r.t. the first occurrence of *x_n_* and not *x_In_*.

#### Definition 1.1

In *G*, a subset *I* ⊆ *V* of nodes is Feedback Vertex Set (FVS) if and only if removal of set *G \ I* leaves a graph without directed cycles. An FVS is minimal if it does not contain a proper subset that is an FVS itself. For simplicity, in this paper, we will consider all the FVSes to be minimal.

#### Definition 1.2

In a dynamic system, a subset *J* ⊆ *V* of nodes is a set of determining nodes if and only if two solutions satisfy 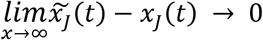 whenever 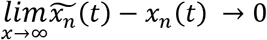 for all components *n* ∈ *J* ⊆ *V*.

In Fielder *et al*. and Mochizuki *et al*. these two definitions were proven to be equivalent for dynamics in a network^5,6^. Therefore, observation of the long-term dynamics of the FVS is sufficient to identify all possible attractors of an entire system. Controlling the dynamics of the FVS 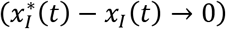 is sufficient to drive the dynamics *x*(*t)* of a whole system to converge on one of any attractors *x*\*(*t*).

The minimal Feedback Vertex Set problem is a well-known NP-hard problem. Many algorithms have been developed to find the near-minimum FVS. Based on the implementation in Zañudo *et al.^76^*, we use a simulated annealing local search approach, SA-FVSP, originally described in Galinier *et al.^55^*. SA-FVSP has been shown to outperform the greedy adaptive search procedure^77^. A network may have multiple FVSes depending on the size and structure, but each FVS has the same capabilities for controlling cell fates.

### Simulating Perturbations on FVS Control Nodes

After identifying the FVS control nodes for virtual screenings, combinations of perturbations (overexpression/upregulation, knockouts/downregulation, or no change) to an FVS node’s activity are generated. NETISCE generates 3*^n^* combinations of control nodes perturbations, where n is the number of FVS control nodes.

The initial state of a node not contained in the FVS or an FVS node whose perturbation is “no change” is set to the normalized expression value of the experimental sample(s) for the selected undesired phenotype.

To simulate the perturbations on FVS control nodes, we modified the SFA algorithm to override the activity of perturbed control nodes. Specifically, the values of the perturbed FVS control nodes are fixed and unaffected by the incoming signal flow. The fixed state *p* of an upregulated (downregulated) FVS control nodes *i* is defined as:

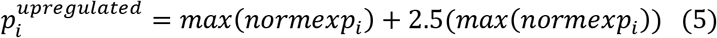

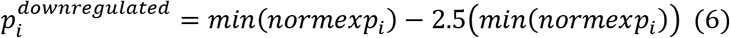

Where *max*(*normexp_i_*) and *min*(*normexp_i_*) are the maximum and minimum normalized expression values of *i* across the experimental samples of the undesired phenotype, respectively. These equations are also used when gain-of or loss-of-function information from mutational data is supplied to NETISCE. For example, the value of a node representing a gene with a gain-of-function mutation is fixed to the corresponding 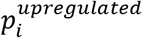 based on the normalized gene expression data.

### Classification of Perturbations on FVS Control Nodes

To systematically identify which perturbations on the FVS control nodes shifted the system away from the attractor associated with the undesired phenotype and towards the desired phenotype, we filter the resultant attractors with two criteria. Criterion 1 considers a combination of perturbations on FVS control nodes successful if the perturbation’s corresponding attractor is classified to the cluster associated with the desired phenotype by at least two of three machine learning classification algorithms. The three classification algorithms considered in NETISCE are Naïve Bayes, Support Vector Machines, and Random Forest classifiers. Naïve Bayes and Support Vector Machines are well suited for high dimensional datasets^78,79^, while Random Forest Classifiers improve predictive accuracy and reduce over-fitting^80^. For each algorithm, the training data is the set of attractors generated from the provided normalized expression data and the randomly generated initial states and their associated clusters identified in the k-means clustering step. For each attractor, the entire vector of all network nodes is supplied. Lastly, we perform a Feature Importance Analysis. We examine the important features (network nodes and their associated steady-state values) used by each algorithm to classify the combinations of perturbations on FVS control nodes. We extract the top 10% of ranked features for each algorithm^81^.

Criterion 2 focuses on the steady-state expression values of the internal-marker nodes. In this second criterion, the attractor obtained after simulating the system under the studied combination of perturbations on FVS control nodes must have at least 90% of the steady-state expression values of its internal-marker nodes within the expected gene expression value ranges of the desired attractors. This ensures that beyond the machine learning classification based on the entire attractor, the known biological internal-marker nodes have the expected values of the desired phenotype. By default, NETISCE is set to strict filtering criteria, where the steady-state expression values of the attractor produced by the control node perturbation must be within the range of the desired phenotype expression values. For example, consider an internal-marker node whose steady-state expression values in the attractors associated with the desired phenotype is 2.0. The steady-state expression values in the attractors associated with the undesired phenotype are 1.0. For a perturbation on the FVS control nodes to pass the filtering criterion, the steady-state expression value of the internal-marker node must be greater than 2.0. Alternatively, the user can select a more relaxed filtering threshold. In this case, for the example described above, a perturbation on the FVS control nodes would pass the filtering criterion if the steady-state expression value of the internal-marker node is greater than 1.0. All the perturbations to the FVS control nodes that pass both filtering criteria are considered to successfully shift the initial state to an attractor associated with the desired phenotype. If NETISCE is run with replicates for the undesired phenotype, then a perturbation on FVS nodes must pass the first filtering criteria on all replicates. All replicates are individually analyzed in the second filtering criterion, and NETISCE produces a separate list of perturbations that pass the criterion for each replicate. In our pluripotent stem cell example that contained three replicates, the perturbations on FVS control nodes that passed both criteria in the replicates were identical. However, users familiar with their data may be interested in perturbations that only work for a subset of replicates.

### Data for Developmental, Stem Cell, and Cancer Biology Validations

#### Cell fate specification in the ascidian embryo

The network structure was obtained from Kobayashi *et al*. ^55^. Since the focus of this example was to reproduce the experimental results of embryonic cell fate specification using Feedback Vertex Set Control and SFA, we performed computations separate from the NETISCE pipeline but using the essential scripts (see GitHub repository and tutorial). Without available normalized gene expression data for the unperturbed embryo, we performed *in silico* simulations to reproduce the cell fate specification results with SFA. The attractor for an unperturbed embryo was estimated by setting the initial activities of two genes necessary for normal embryonic development, Gata.a and Zic-r.a, to 1, representing an activated state^54^. All other nodes were initialized to 0, representing an initial inactive activity. The attractor estimation function simulated the seven perturbations to the FVS control nodes that induced the seven tissue fates experimentally: (Foxa.A, Foxd, Erk Signaling, Neurog, Tbx6-r.b, and Zic-r.b.). Specifically, in these simulations, Gata.a and Zic-r.a had initial activities set to 1 and all other nodes set to 0. Then, the values of the FVS control nodes were overridden using the 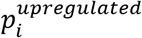 or 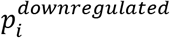 equations. Since there was no gene expression data, for all FVS control genes the 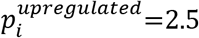 and 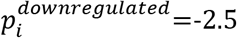. The internal-marker node steady-state expression values for the unperturbed and perturbations on FVS node simulation results can be found in Supplementary Table 1. A perturbation was considered successful in replicating the experimental results if the difference of the steady-state expression values between the specified internal-marker node for the relevant tissue in the attractor generated from a perturbation on FVS control nodes and the attractor generated from the unperturbed initial state was positive (Supplementary Table 4). These values were additionally graphed using radar plots to visualize the respective upregulations for each perturbation (Figure 4).

#### Induced pluripotent stem cell reprogramming from primed to naïve pluripotency

The intracellular signaling network for induced pluripotent stem cell signaling was obtained from Yachie-Kinoshita *et al*.^56^. The normalized expression data for EpiSCs and ESCs were downloaded from the Gene Expression Omnibus (GSE88928)^82^. There were three replicates for each experimental sample. Each replicate was used separately as initial state values to simulate the network, compute their associated steady-states, and perform perturbations on the FVS control nodes. Initially, we selected as internal-marker nodes the four output nodes used in the Boolean Model of Yachie-Kinoshita *et al*.: Oct4, Sox2, Nanog, and EpiTFs. Although these three nodes were also FVS control nodes, they were used as internal-marker nodes to be consistent with the output nodes in the Boolean simulations. To further filter our perturbations, we selected additional internal-marker nodes from gene expression data provided by Yachie-Kinoshita *et al*. ^56^. Based on the gene expression data for the network nodes, there were six genes whose values differed significantly between the ESC and EpiSC states. These included Lefty1, Pitx2, and Esrrb. The three other genes, Tbx3, Gata6, and Klf4, were not included as internal-marker nodes as they were FVS control nodes. Radar plots were used to visualize the perturbations of Klf4 upregulation, Nanog upregulation, and the combined Klf4+Nanog upregulation (Figure 6).

#### Overcoming adaptive resistance to MAPK inhibitory therapy in colorectal cancers

The colorectal cancer (CRC) tumorigenesis signaling network and annotated HT29 mutational profile for network nodes was provided by Park *et al*.^58^. The RNA-seq from untreated HT29 cells was obtained from the Cancer Cell Line Encyclopedia (CCLE)^83^.

In this study of adaptive resistance to MAPKi therapy in CRC, the ultimate therapeutic goal is to decrease proliferation and increase apoptosis in tumor cells. In a method adapted from Beal *et al*.^59^, we verify that CRC tumors’ proliferation and apoptosis signatures are preserved under the SFA simulation of the generic CRC network for patient tumors (Supplementary Text 5).

The network was simulated using as initial conditions the normalized expression and mutational profile of an untreated HT29 as there was no available gene expression data for treated HT29 cells. As annotated in Park *et al*.^58^, PIK3CA and BRAF have gain-of-function mutations, while APC, SMAD4, and TP53 have loss-of-function mutations in HT29 cells. Therefore, the values of these nodes were fixed to the appropriate 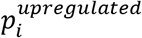 or 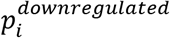 value. To simulate BRAFi (HT29_BRAFi), 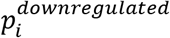 override was applied to the state of BRAF. To simulate BRAFi+EGRFi (HT29_BRAFi+EGRFi), BRAF and EGFR had the appropriate overrides applied using the 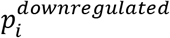 equation. We used the FVS finding algorithm to search for a sufficiently large number of FVS in the CRC network, which identified 68 FVSes of size 14. TP53 was removed from the FVSes since it was already fixed to its 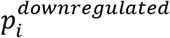 value due to the loss-of-function mutation in HT29 cells (Supplementary Data 1). Based on Feedback Vertex Set control theory^5^, all FVSes of a system can guide the system to any of its natural attractors; therefore, we randomly selected the first FVS identified by the algorithm to perform virtual screenings. For simulating perturbations on FVS control nodes, the system was initialized with the same parameters as the (HT29_BRAFi) simulation, with the additional perturbations to the FVS control nodes.

Control node perturbations were filtered first by the set of 3 internal-marker nodes used by Park *et al*.^58^: CASP3, a marker of apoptosis, CCNE1 (also known as Cyclin E), a proliferation marker, and ERK (also known as MAPK1), a downstream molecule of the MAPK signaling pathway whose activity after BRAFi treatment indicates adaptive resistance. Potential additional internal-marker nodes were selected from downstream signaling elements of MAPK signaling, apoptosis-related, and proliferation-related pathways^58^. These sets were filtered using the internal-marker node checking step of NETISCE to ensure that their steady-state expression values in the attractors associated with MAPK inhibitor therapy and MAPK inhibitor resistance were appropriate based on literature evidence. This resulted in an additional 17 internal-marker nodes: four genes from apoptosis-related pathways (CASP9, DIABLO, MAPKAPK2, PPP2CA), seven genes related to proliferation (CCNB1, CCND1, CDC25A, CDKN1A, CDKN2B, E2F1, RB1), and six genes from MAPK signaling pathways (DUSP1, ELK1, HNF1B, MAPK8, MLK3, RPS6KA1).

##### Evaluating the Robustness to Noise of NETISCE

In COPASI, we simulated the DE models using the Time Course function for the undesired and desired phenotypes. We additionally simulate the time course for the undesired initial condition when a perturbation on FVS nodes is applied to ensure the system still arrives at the desired attractor. We injected seven levels of noise (1%, 5%, 10%, 20%, 30%, 40%, 50%) in the undesired and desired initial conditions using the Random Distribution item in the COPASI’s Parameter Scan function. For each node with a nonzero initial concentration, the noisy initial condition was generated using a normal distribution, where the mean was the initial state of the node, and the standard deviation was .1, .5, .10, .20, .30, .40, or .50, to simulate 1%, 5%, 10%, 20%, 30%, 40%, or 50% noise, respectively. We generated 1,000 initial states for each noise level for the desired initial and undesired initial states.

In COPASI, von Dassow’s Drosophila Segment Polarity Gene model simulations were computed using the deterministic LSODA Solver for 500 seconds when a steady-state was reached. The values of the FVS control nodes were set to the values determined by *Zanudo et al*. (Supplementary Table 1). The SDE model was extracted from Zhou *et al*. and simulated using the SDE solver. To implement the time-delay perturbations of MafA, Pdx1, Ngn3, Pax4 overexpression, and Ptf1a knockout in exocrine cells, we used the Event function to increase the production or degradation rates as performed in *Zhou et al*.

After completing the simulations in COPASI and generating sets of noisy initial conditions, we tested the ability of NETISCE to correctly identify that the specified perturbations on FVS control nodes can shift the undesired initial state away from the undesired attractor and towards the desired attractor. It is unlikely that an experiment contains 1,000 biological replicates in real circumstances. Therefore, we generated 1,000 subsets of three wild-type and three unpatterned initial states at each noise level, analogous to an experiment with three biological replicates of the wild-type phenotype and three biological replicates of the unpatterned phenotype. For each set of 1,000 initial states at each noise level, we run NETISCE with default settings.

### NETISCE Implementation

The main computational scripts of our pipeline are written in Python, utilizing the extensively optimized machine learning algorithms of the Scikit Learn package^81^. Scripts for analyzing the internal-marker node values are written in R. NETISCE is implemented as a Nextflow workflow^84^. Nextflow is a state-of-the-art workflow manager tool that is language agnostic and designed for parallel processing as a dataflow manager. Checkpoints are implemented for the user to investigate any possible errors or make changes to the run configuration. The code can easily be resumed without having to re-run all computations. We provide Nextflow pipelines for local machine use and high-performance cluster implementations. We also provide NETISCE within a Docker container to further enhance the reproducibility of NETISCE simulations^85^. In addition to the command-line tool, our pipeline is available through the Galaxy Project. This cloud-based open-source tool requires little to no programming experience for biological analysis and workflows^86^.

## Data availability

The NETISCE’s Nextflow pipeline version, the Docker image documentation, and data are available at https://github.com/veraliconaresearchgroup/netisce. The installation, tutorials, information for installing the Galaxy Project version of NETISCE, and walkthroughs for reproducing the above results are found at https://veraliconaresearchgroup.github.io/Netisce/.

## Supplementary Files

Supplementary Information 1: supplementary-file.pdf

Supplementary Data 1: supplementary-data1.xlsx

## Acknowledgments

We thank Pedro Mendes and Ion Moraru for their insightful discussions and feedback in the methodology and validation of NETISCE. We additionally thank Pedro Mendes for his help with the COPASI simulations of the differential equations models. S.B. was supported by the UConn Health Research Program; A.P. and M.S. worked on this project under the UConn High School Mentored Research Program.

## Author contributions

P.V-L. and L.M. worked in the conceptualization and planning of this work. L.M, M.S., and A.P. performed and analyzed the validation simulations. L.M, S.B., and M.S. participated in the coding and implementation of NETISCE in Nextflow and Galaxy Project. L.M. and P.V-L. drafted the manuscript. All authors read and approved the manuscript.

