## Supplementary Information for "NETISCE: A Network-Based Tool for Cell Fate Reprogramming"

**List of Supplementary Information**


| | |
|---|---|
| **Supplementary Text 1:** | Overview of Network-Based and Dynamical Systems-Based Approaches to Cell Fate reprogramming |
| **Supplementary Figure 1** | Dynamical Systems-Based Approaches to Cell Fate reprogramming |
| **Supplementary Text 2:** | Validation of NETISCE in *in silico* studies of cell fate reprogramming using the FVS Control in ODE and Boolean Models |
| **Supplementary Table 1:** | Summary ODE simulations of Drosophila segment polarity gene network |
| **Supplementary Figure 2:** | Boolean Drosophila segment polarity gene network |
| **Supplementary Table 2:** | Summary Boolean simulations of Drosophila segment polarity gene network |
| **Supplementary Table 3:** | Steady-state values for the study of the cell fate specification in Ascidian embryos |
| **Supplementary Table 4:** | Differences between the values of the internal-marker nodes in the attractors generated from the seven experimentally verified perturbations on the FVS control nodes and the values in the unperturbed state for cell fate specification in Ascidian embryos. |
| **Supplementary Table 5:** | Table of the fifteen perturbations on FVS control nodes that passed both filtering criteria for induced pluripotent stem cell reprogramming from primed to naïve pluripotency simulations when using seven internal-marker nodes. |
| **Supplementary Figure 3:** | Bar plots for each FVS control node in the fifteen perturbations that passed both filtering criteria for induced pluripotent stem cell reprogramming. |
| **Supplementary Text 3:** | Feature Importance Analysis in iPSC Reprogramming Machine Learning Classification |
| **Supplementary Table 6:** | Top 10% of Features of iPSC Reprogramming Machine Learning Classification Algorithms |
| **Supplementary Text 4:** | Network Revision Based on NETISCE Simulations |
| **Supplementary Data 1:** | The 68 FVSes in the colorectal cancer signaling network. |
| **Supplementary Text 5:** | Methods and results of Validation of Colorectal Cancer Network and SFA Simulation. |

**Supplementary Figure 4:** Validation of Colorectal Cancer (CRC) Signaling Network and SFA simulation

**Supplementary Text 6:** Feature Importance Analysis in CRC MAPKi Resistance Reprogramming Machine Learning Classification

**Supplementary Table 7:** Top 10% of Features of CRC MAPKi Resistance Reprogramming Machine Learning Classification Algorithms

**Supplementary Figure 5:** Bar plots for each of the FVS control nodes in the 1,266 perturbations that passed both filtering criteria in the colorectal cancer adaptive resistance reversion simulations

**Supplementary Figure 6:** COPASI simulations of von Dassow's Drosophila Segment Polarity Gene ODE model

**Supplementary Table 8:** Summary of the NETISCE simulations of control in von Dassow's Drosophila Segment Polarity Gene model with noisy initial states

**Supplementary Figure 7:** Reprogramming Pancreatic Exocrine Cells to Beta-Cells

**Supplementary Text 7:** Reprogramming Pancreatic Exocrine Cells to Beta-Cells in NETISCE

**Supplementary Table 9:** Summary of the NETISCE simulations in Pancreatic Cell Fate Specification model with noisy initial states.

The raw data files where the tables and figures were generated, and the code used to produce them, can be found in the GitHub repository (https://github.com/veraliconaresearchgroup/netisce) and the NETISCE manual website (https://veraliconaresearchgroup.github.io/Netisce/).

**Overview of Network-Based and Dynamical Systems-Based Approaches to Cell Fate reprogramming**

The growth of computational methods for constructing biological interaction networks (e.g., gene regulatory and signaling networks) from genome-wide expression data[1–3] has led to the development of cell reprogramming methods within the framework of network biology. Network-based approaches such as CellNet and ANANSE use gene expression profiles to construct gene regulatory networks (GRNs) and evaluate reprogramming experiments[4,5]. These algorithms score cellular reprogramming experiments by analyzing the extent to which a reprogrammed cell establishes the desired phenotype's GRN and suggest transcription factors to induce the desired reprogramming. For example, CellNet was used to determine the reprogramming efficiency of fibroblasts to hepatocyte-like cells (iHeps) by comparing GRNs of fibroblasts, hepatocytes, and the reprogrammed fibroblasts[6]. CellNet analysis revealed that the reprogrammed cells did not exhibit the hepatocyte cell identity and suggested that downregulation of the transcription factor Cdx2 could drive these cells towards the hepatocyte-like state.

However, a significant limitation of static network-based approaches is that cell reprogramming does not necessarily arise only from the network topology. A systems dynamics contain essential information for cellular reprogramming that is impossible to capture in a purely static view of cell phenotypes[7]. The dynamical systems framework can be used to derive a quantitative understanding of cellular reprogramming[8–10], where each cell fate is a stable state (attractor) shaped by the architecture and the dynamics of its regulatory network[10–13] (Supplementary Figure 1a). Analyzing a mathematical model's phase space of a system is an established dynamical systems-based framework for studying cell fate differentiation, tumorigenesis, and ultimately, the reversion of these processes[14–19]. For example, phase space analysis of an Ordinary Differential Equations model (ODE) of pancreatic cell fate regulation revealed branching points in pancreatic cell differentiation[18]. Then, transcription factor perturbations were simulated to study their effect on pancreatic cell differentiation trajectories in the system. These simulations uncovered that the mechanism behind experimentally-observed exocrine pancreatic cell to beta-cell reprogramming required triggering the cell to pass through one of the branching points. Additional analysis revealed that sequentially perturbating the relevant transcription factors could improve reprogramming efficiency.

In discrete modeling frameworks, such as Boolean networks modeling, the phase space can be studied by analyzing the attractor landscape (Supplementary Figure 1b). For instance, a quantitative evaluation of the attractor landscape was performed to study colorectal cancer tumorigenesis and its reversion in the work of Cho *et al.* and Kim *et al*. [15,16]. In these works, the attractor landscape analysis was performed on a Boolean network model of colorectal cancer, and regions of the attractor landscape were associated with normal-like or malignant phenotypes[15,16]. Then, when single and double perturbations for all nodes in the network were simulated, the attractors were evaluated to determine if the perturbation shifted the system to the region associated with the normal-like phenotype, indicating potential targets for tumor reversion. In combination with a control theory-based approach, this attractor landscape analysis was recently applied in a Boolean Model of Basal-Like Breast Cancer Reprogramming, where targets that can induce basal-like breast cancer reprogramming and endocrine therapy sensitivity were identified[20].

One powerful method for identifying combinations of targets to guide induced pluripotent stem cell (iPSC) reprogramming is IRENE[21], which combines both network-based and dynamical systems-based approaches. IRENE takes a multi-omics approach to constructing GRNs for various cell types, incorporating data such as transcriptomic, epigenetic, and enhancer-promoter interaction profiles. Next, the method infers logical rules for the models using ChIP-seq and protein-protein interaction data and performs stochastic simulations on the iPSC network. Finally, based on the probabilities generated from the simulations, IRENE identifies combinations of transcription factors that, when perturbed, activate the target cell type GRN by considering the number of epigenetic changes needed to shift the enhancer/promoter landscape to the target cell type. This tool was shown to increase reprogramming efficiencies for iPSC differentiation to melanocytes and natural killer cells and generated reprogramming targets for iPSCs to mammary epithelial cells.

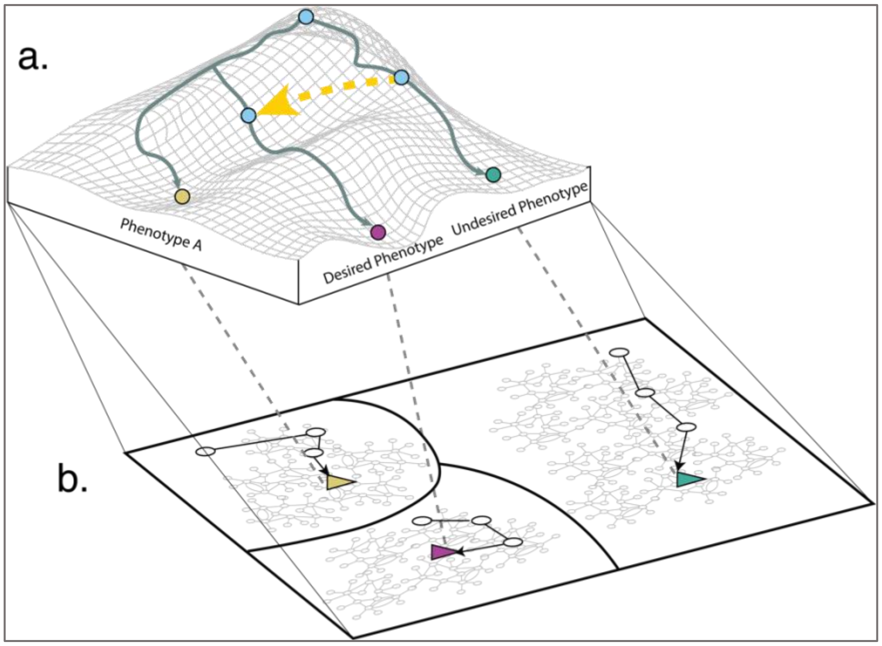


**Supplementary Fig.1 Dynamical Systems and Control Theory View of Cell Fate Reprogramming. a** The Waddington's epigenetic landscape metaphor to visualize cell fate reprogramming: a cell evolves until it reaches a long-term steady-state or attractor (green circle) that can be associated with an observable phenotype (*e.g.,* undesired phenotype). Cell fate reprogramming consists of shifting (yellow dashed arrow) the state of the cell from one trajectory towards another, which attractor state (purple circle) is associated with the desired phenotype. **b** Alluding to the Boolean network's framework, the attractor landscape of a system represents all potential states of a system and its attractors. For a given attractor, its basin of attraction is the set of initial conditions leading to that attractor, as represented by the three different sections in the figure. The attractors and their basins of attraction can be associated with cell phenotypes.

**Validation of NETISCE in *in silico* Studies of Cell Fate Reprogramming using the FVS Control in ODE and Boolean Models**

In these application examples, we compare the ability of NETISCE to correctly simulate perturbations on FVS control nodes to guide a system from an undesired initial state to the desired state for two in silico studies of cell fate reprogramming using the FVS control in Boolean and ODE models of Drosophila embryonic patterning.

To assess the FVS control approach, Zanudo *et al*.[22] used the Drosophila Melanogaster segment polarity network that guides gene expression during embryonic development. This study aimed to show the ability of the FVS control to steer the dynamics of a system towards any natural attractor in a validated model. With an established ODE[23] and Boolean Model[24] of the segment polarity gene network, the authors showed that fixing the values of the FVS nodes to their state in the correctly patterned attractor (the wild-type attractor) was sufficient to guide the system to the patterned attractor irrespective of the initial state of all other network nodes.

The continuous ODE model of von Dassow *et al*.[23] represents each cell as a hexagon with six relevant cell-to-cell boundaries (Supplementary Figure 6a). It includes 136 nodes representing mRNAs and proteins: four source nodes and 24 sink nodes, and 488 edges representing transcriptional regulation, translation, and protein-protein interactions. The nodes are characterized by continuous concentrations, whose rate of change is described by ordinary differential equations (ODE) involving Hill functions for gene regulation and mass action kinetics for protein-level processes and using 48 kinetic parameters. From Zanudo *et al*.[22], we obtained the initial conditions that produce the wild-type attractor, the unpatterned attractor, and the precise combination of perturbations on the 48 FVS nodes that shifts the unpatterned initial state from the trajectory of the unpatterned attractor towards the wild-type patterned attractor (Supplementary Table 1). In addition, we considered as internal-marker nodes those used by Zanudo *et al*. to characterize the wild-type, unpatterned, and the controlled unpatterned-to-wild-type attractors: *wg1*(representing the wingless gene in the first cell) and *en2* (engrailed in the second cell).

**Supplementary Table 1:** Summary of initial conditions that yield the wild-type or unpatterned steady state, and the configuration of perturbations on the FVS control nodes to shift the system from the unpatterned attractor towards the wild-type attractor in the ODE model of Drosophila segment polarity genes, reproduced from Zanudo *et al*.[22].

| **Initial condition leading to wild-type steady state** | **Initial condition leading to unpatterned steady state** | **Perturbations on FVS control nodes to shift the unpatterned initial state to converge to the wild-type attractor (identified by Zanudo *et al*.[22])** |
|---|---|---|
| wg /IWG in the first cell is at maximal concentration = 1 | Intra-cellular nodes have a concentration of 0.05 in the first and third cell | overexpression: 1 EWG 0, 1 EWG 2, 1 EWG 4, 1 IWG |
| en /EN in the second cell has concentration = 1 | Intra-cellular nodes have a concentration of 0.15 in the second and fourth (zeroth) cell | knock-out: 0 CN, 0 EWG 0, 0 EWG 2, 0 EWG 4, 0 HH 0, 0 HH 2, 0 HH 4, 0 IWG, 0 PTC 0, 0 PTC 2, 0 PTC 4, 0 ptc, 1 CI, 1 CN, 1 HH 0, 1 HH 2, 1 HH 4, 1 PTC 0, 1 PTC 2, 1 PTC 4, 2 CI, 2 CN, 2 EWG 0, 2 EWG 2, 2 EWG 4, 2 HH 0, 2 HH 2, 2 HH 4, 2 IWG, 2 PTC 0, 2 PTC 2, 2 PTC 4, 3 CN, 3 EWG 0, 3 EWG 2, 3 EWG 4, 3 HH 0, 3 HH 2, 3 HH 4, 3 IWG, 3 PTC 0, 3 PTC 2, 3 PTC 4, 3 ptc |
| Source nodes (B) are fixed at 0.4 in each cell | Membrane-localized nodes have concentration of 0.15 for even-numbered sides in every cell | |
| Membrane-localized nodes have a concentration of 0.05 for odd-numbered sides in every cell | | |

We tested the ability of NETISCE to correctly identify that the perturbations on FVS control nodes can shift the unpatterned initial state away from the unpatterned attractor and towards the wild-type attractor. Therefore, we modified NETISCE to only consider the perturbation on FVS control nodes specified by Zanudo *et al*. Otherwise, NETISCE was run with default settings.

K-means clustering was performed on the attractors generated from the wild-type and unpatterned initial states and the 100,000 randomly generated initial states. According to the elbow metric, the k-means optimality metrics returned an optimal k of k=3, and k=2 by the silhouette metric. NETISCE proceeded with k-means clustering with k=2. One cluster contained the attractors generated from the wild-type initial states, and one cluster contained the attractors generated from the unpatterned initial states. The specified perturbation on FVS control nodes passed both filtering criteria and thus successfully shifted from the unpatterned to wild-type cell fate.

The Boolean model of Albert and Othmer[24] represents four cells as a repeating unit to reproduce wild-type stable cell patterning. This model contains 56 nodes and 144 edges, representing the four cells as hexagons with two cell-to-cell boundaries (Supplementary Figure 2). In this model, Zanudo *et al.* focused on hedgehog expression in cell 2 (hh2) and engrailed in cell 2 (eg2), which we consider internal-marker nodes.

Zanudo *et al.* identified two initial conditions for network nodes that yielded either the wild-type patterned steady state or the unpatterned steady state. They identified an FVS consisting of the following nodes: 0_PTC, 0_hh, 0_wg, 1_PTC, 1_wg, 2_PTC, 2_hh, 2_wg, 3_PTC, and 3_wg. Using the Boolean Model, Zanudo *et al.* simulated the results of FVS and source node perturbations. From any arbitrary initial condition, including the unpatterned initial state, fixing the state of the FVS and source nodes to their values at the wild-type steady state resulted in the wild-type steady state (Supplementary Table 6).

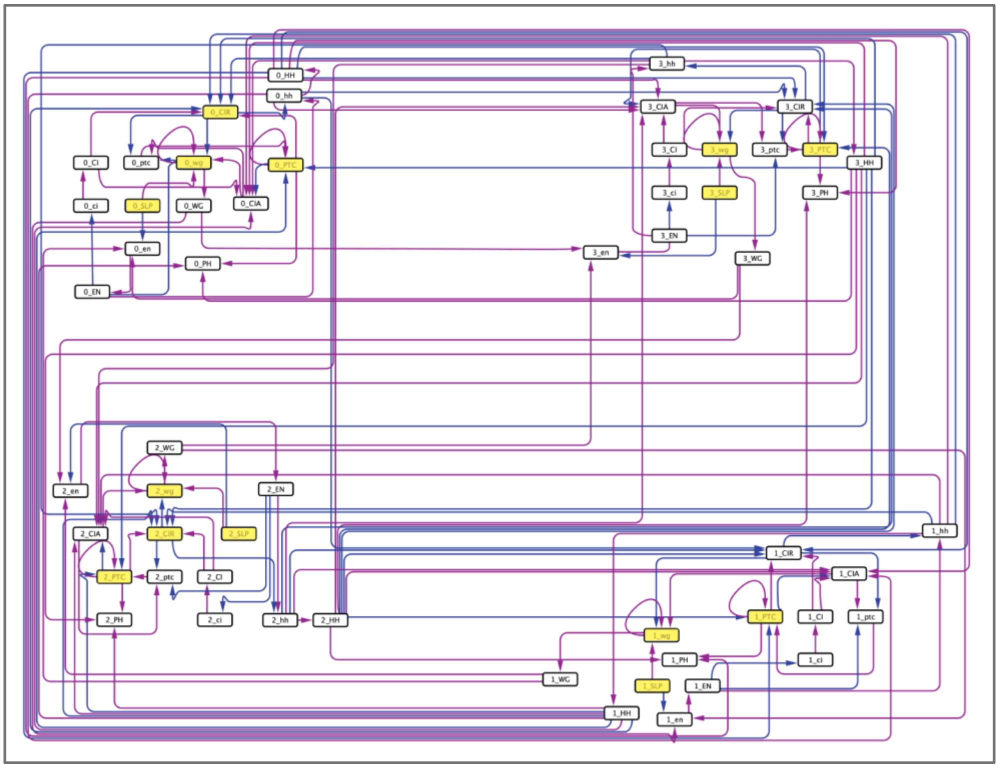

**Supplementary Figure 2: Drosophila Segment Polarity GRN.** Purple edges indicate activating interactions, while blue edges indicate inhibitory interactions. Nodes highlighted in yellow are members of the FVS and source node set explored in Zanudo *et al*.[22].

**Supplementary Table 2:** Summary of the Boolean simulations of the Drosophila segment polarity gene network reproduced from Zanudo *et al*.[22].

| Steady state | Initial conditions | Steady-state values |
|---|---|---|
| Wild-type | ON: | $hh_2$:ON<br>$en_2$: ON |
| Unpatterned | OFF: all nodes | $hh_2$: OFF<br>$en_2$: OFF |
| Unpatterned initial state + FVS controlled | ON: : 0_PTC, 0_hh, 0_wg, 1_PTC, 1_wg, 2_PTC, 2_hh, 2_wg, 3_PTC,3_wg | $hh_2$: ON<br>$en_2$: ON |

We again tested the ability of NETISCE to correctly identify that the perturbations on FVS control nodes can shift the unpatterned initial state away from the unpatterned attractor and towards the wild-type attractor. Like the ODE model, the k-means optimality metrics returned an optimal k of k=3, according to the elbow metric, and k=2 by the silhouette metric. NETISCE proceeded with k-means clustering with k=2, where one cluster contained the attractors generated from the wild-type initial state, and one cluster contained the attractors generated from the unpatterned initial state. The specified perturbation on FVS control nodes passed both filtering criteria and thus successfully shifted from the unpatterned to wild-type cell fate. These studies show that our attractor landscape estimation and SFA simulation framework in NETISCE successfully reproduces the *in silico* cell fate reprogramming studies using the FVS control in ODE and Boolean modeling frameworks.

**Supplementary Table 3:** Steady-state values for the study in Ascidian embryos' cell fate specification. The steady-state values produced from SFA simulations for the seven internal-marker nodes in the attractor from an unperturbed simulation and the attractors generated from the seven perturbations on the FVS control nodes that were experimentally verified in Kobayashi *et al.*[25]. The first column lists the name of the simulation: either unperturbed or the combination of perturbation to the FVS control nodes, where an uppercase letter indicates overexpression, and a lowercase letter indicates knockout. The column names are the internal-marker nodes for each tissue fate: Alp=endoderm, Bco=Brain, Celf.a=pan-neural, Epi1=epidermis, Fli/Erg.a=mesenchyme, Myl=muscle, Noto1=notochord.

| name | Alp | Bco | Celf3.a | Epi1 | Fli.Erg.a | Myl | Noto1 |
|---|---|---|---|---|---|---|---|
| unperturbed | -3.41E-05 | 1.87E-03 | 1.87E-03 | 2.18E-02 | 1.40E-03 | 8.98E-04 | 4.03E-04 |
| Adentz (Endoderm perturbation) | 1.18E-01 | -4.25E-01 | -4.25E-01 | 2.71E-01 | -1.84E-01 | -1.40E-01 | 1.17E-01 |
| adentZ (brain+pan-neural perturbation) | -3.85E-01 | 4.25E-01 | 4.25E-01 | 2.71E-01 | -2.37E-01 | -4.16E-03 | 1.12E-01 |
| adeNtz (pan-neural perturbation) | -5.24E-01 | -4.25E-01 | -4.25E-01 | 2.71E-01 | -3.00E-01 | -1.06E-01 | 2.41E-01 |
| adEntZ (mesenchyme perturbation) | -2.63E-01 | 4.25E-01 | 4.25E-01 | -1.56E-01 | 2.84E-01 | -8.71E-02 | -5.26E-03 |
| adentz (epidermis perturbation) | -5.24E-01 | -4.25E-01 | -4.25E-01 | 2.71E-01 | -3.00E-01 | -1.24E-01 | 2.35E-01 |
| adenTz (muscle perturbation) | -5.24E-01 | -4.25E-01 | -4.25E-01 | 2.71E-01 | -3.00E-01 | 8.37E-02 | 2.59E-01 |
| aDentz (notochord perturbation) | -3.79E-01 | -4.25E-01 | -4.25E-01 | 1.99E-01 | -3.97E-01 | -1.22E-01 | 9.43E-02 |

**Supplementary Table 4:** The differences between the internal-marker nodes' values in the attractors generated from the seven experimentally verified perturbations on the FVS control nodes and the values in the unperturbed state for cell fate specification in Ascidian embryos. The values were calculated by subtracting the steady-state value in the unperturbed attractor from the steady-state value produced from the perturbation on the FVS control nodes. Positive values indicate that the specified gene is upregulated in the attractor produced from the perturbation on FVS control nodes compared to the unperturbed attractor. For a simulation of the perturbations on FVS control nodes to be considered successful, the steady-state gene expression value of the internal-marker node must be greater in the attractor produced by the perturbation on the FVS control nodes than the steady-state gene expression value in the attractor associated with the unperturbed state. Yellow-highlighted cells denote perturbations that produced the desired internal-marker node upregulation. The column names are the internal-marker nodes for each tissue fate: Alp=endoderm, Bco=Brain, Celf.a=pan-neural, Epi1=epidermis, Fli/Erg.a=mesenchyme, Myl=muscle, Noto1=notochord. The row names are the abbreviations of the combinations of perturbations on FVS control nodes (A/a=Foxa.a, D/d=Foxd, E/e= Erk signaling, N/n= Neurog, T/t=Tbx6-r.b, Z/z=Zic-r.b, uppercase indicates overexpression, lowercase indicates knockout). The tissue that was induced experimentally is listed in parentheses.

| **name** | **Alp** | **Bco** | **Celf3.a** | **Epi1** | **Fli.Erg.a** | **Myl** | **Noto1** |
|---|---|---|---|---|---|---|---|
| Adentz (experimentally induced endoderm) | 0.118 | -0.427 | -0.427 | 0.249 | -0.185 | -0.141 | 0.117 |
| adentZ (experimentally induced brain+pan-neural) | -0.385 | 0.423 | 0.423 | 0.249 | -0.238 | -0.005 | 0.112 |
| adeNtz (experimentally induced pan-neural) | -0.524 | -0.427 | -0.427 | 0.249 | -0.302 | -0.107 | 0.24 |
| adEntZ (experimentally induced mesenchyme) | -0.263 | 0.423 | 0.423 | -0.178 | 0.283 | -0.088 | -0.006 |
| adentz (experimentally induced epidermis) | -0.524 | -0.427 | -0.427 | 0.249 | -0.302 | -0.125 | 0.235 |
| adenTz (experimentally induced muscle) | -0.524 | -0.427 | -0.427 | 0.249 | -0.302 | 0.083 | 0.259 |
| aDentz (experimentally induced notochord) | -0.379 | -0.427 | -0.427 | 0.177 | -0.398 | -0.123 | 0.094 |

**Supplementary Table 5:** Table of the fifteen perturbations on FVS control nodes that passed both filtering criteria for induced pluripotent stem cell reprogramming from primed to naïve pluripotency simulations when using seven internal-marker nodes. The first column contains the perturbation ID for the fifteen perturbations out of the 729 total possible perturbations. Columns two to seven contain the specified perturbation to each FVS control node: “up” denotes overexpression, “down” denotes knockouts, and blank cells represent no change to the node’s activity. The last three columns contain information about the number of overexpression and knockout perturbations and the total number of nodes perturbed in the considered FVS set.

| name | **Sox2** | **Nanog** | **Gata6** | **Tbx3** | **Oct4** | **Klf4** | **# overexpression** | **# knockouts** | **total** |
|---|---|---|---|---|---|---|---|---|---|
| pert_445 | | up | | | | | 1 | 0 | 1 |
| pert_418 | | up | down | | | | 1 | 1 | 2 |
| pert_436 | | up | | down | | | 1 | 1 | 2 |
| pert_446 | | up | | | | up | 2 | 0 | 2 |
| pert_454 | | up | | up | | | 2 | 0 | 2 |
| pert_409 | | up | down | down | | | 1 | 2 | 3 |
| pert_419 | | up | down | | | up | 2 | 1 | 3 |
| pert_427 | | up | down | up | | | 2 | 1 | 3 |
| pert_437 | | up | | down | | up | 2 | 1 | 3 |
| pert_455 | | up | | up | | up | 3 | 0 | 3 |
| pert_473 | | up | up | | | up | 3 | 0 | 3 |
| pert_410 | | up | down | down | | up | 2 | 2 | 4 |
| pert_428 | | up | down | up | | up | 3 | 1 | 4 |
| pert_464 | | up | up | down | | up | 3 | 1 | 4 |
| pert_482 | | up | up | up | | up | 4 | 0 | 4 |

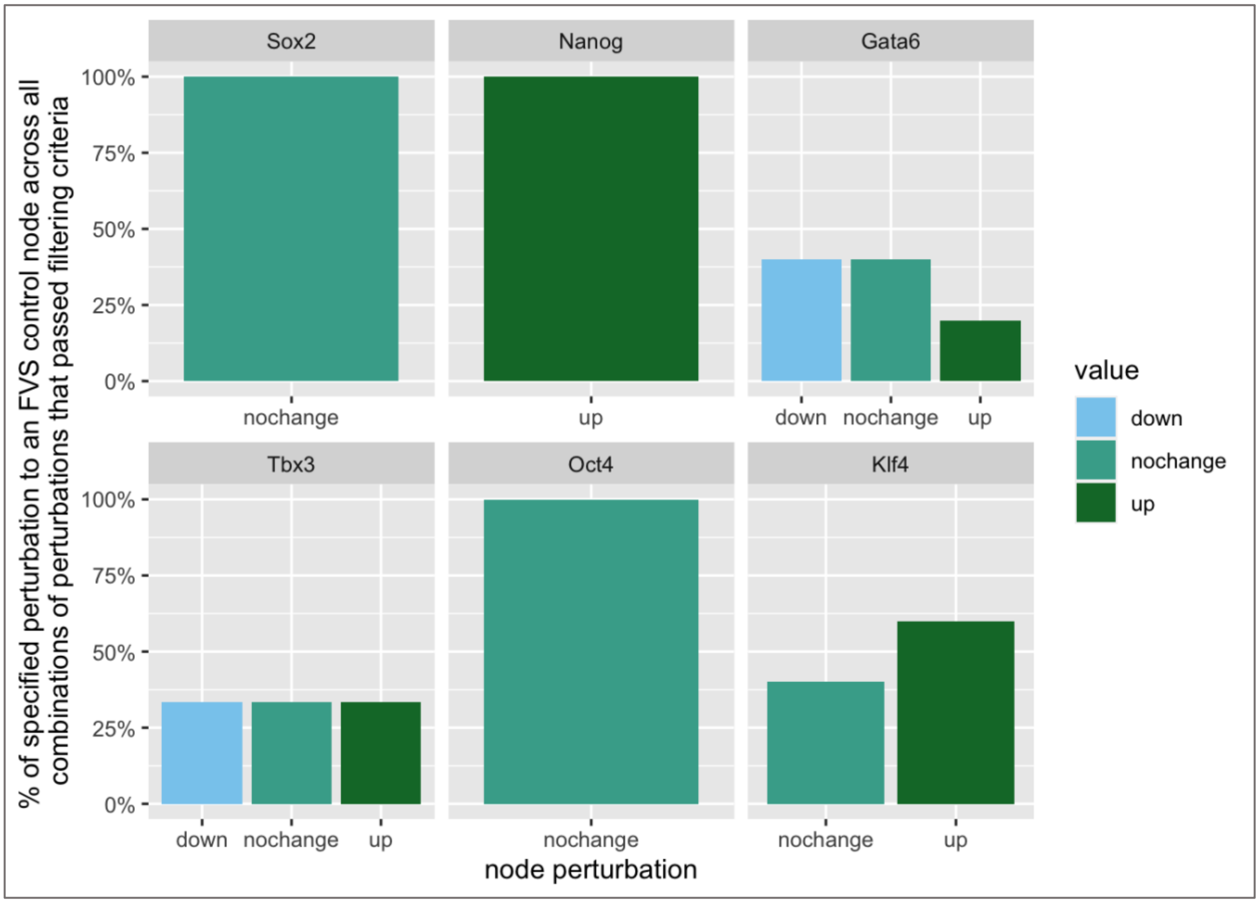


**Supplementary Figure 3:** Bar plots for each FVS control node in the fifteen perturbations that passed both filtering criteria for induced pluripotent stem cell reprogramming from primed to naïve pluripotency simulations when seven internal-marker nodes are used for Criterion 2. These plots describe the percentages of overexpression knockout perturbations or no change to activity for an FVS control node across the fifteen perturbation sets. The x-axis contains the denoted perturbation (down=knockout, up=overexpression, or no change). The y-axis denotes the percentage of knockout, overexpression perturbations, or no change to that node across all fifteen sets of perturbations on FVS control nodes.

### Feature Importance Analysis in iPSC Reprogramming Machine Learning Classification

NETISCE employs three machine learning classification algorithms to assess whether a given set of perturbations on an FVS set does shift the system from a given initial state to the desired cluster of attractors associated with the desired phenotype. We investigated how each algorithm classifies these perturbation sets on FVS control nodes by performing a Feature Importance Analysis for the Random Forest, Naive Bayes, and SVM classifiers (see Methods). In other words, we wanted to examine which network nodes and their steady-state expression values are critical determinants for classifying the attractors generated from perturbations on FVS control nodes. We obtained each machine learning classification method's top 10 percent ranked features (Supplementary Table 6). We observe that the top features across all three classification algorithms are mutually exclusive. In addition, two of the highest-scoring nodes for SVM classification, Klf4, and Tbx3, are FVS control nodes. Note that only one node, Dusp6 had a non-zero importance score for Naive Bayes classification. Interestingly, zero of the attractors generated from combinations of perturbations on FVS control nodes were classified to the ESC cluster by Naive Bayes (all 375 attractors that passed criterion 1 were classified to the ESC associated cluster by both SVM and Random Forest classifiers).

**Supplementary Table 6**: Top 10% of important features for SVM, Naive Bayes, and Random Forest classification algorithms in iPSC network simulations.

| SVM | Naive Bayes | Random Forest |
|---|---|---|
| Klf4 | Dusp6 | Pecam1 |
| Klf2 | | Lrh1 |
| Tbx3 | | Fgf4 |

### Network Revision Based on NETISCE Simulations

Based on the Naive Bayes' Feature Importance Analysis and classification results, we investigated if NETISCE was not correctly predicting the value of Dusp6 at the steady state. We observed that the gene expression value of Dusp6 was greater in the attractors generated from the EpiSC initial states than in the attractors generated from the ESC samples. This observation disagreed with the literature, which states that high levels of Dusp6 maintain pluripotency[26].

Additionally, if Dusp6 were considered a marker node, the perturbations on FVS control nodes would fail to pass criterion 2 because the steady-state expression values of Dusp6 would not shift to the expression values in the attractors generated from the ESC initial states. The disagreement between literature information and the inability to shift towards the desired state indicated that the network might be missing information to simulate Dusp6 accurately. We used the INDRA[27] database tool to search for literature and database information to determine if there was missing edge information between other network nodes and Dusp6. INDRA identified an inhibitory edge between Pitx2 and Dusp6. This edge was added to the network, and NETISCE simulations were re-computed. Indeed, the steady-state expression value of Dusp6 agreed with the literature expectation and was greater in the attractors generated from the ESC initial state data than the attractors generated from the EpiSC initial state data. With the modified network, 390 attractors generated from perturbations on FVS control nodes were classified to the cluster associated with the ESC phenotype by both SVM and Random Forest Classifiers. The Naive Bayes algorithm did not classify any attractors to the ESC-associated cluster. The 390 attractors comprised the same 372 attractors generated by combinations of perturbations on FVS control nodes in the unmodified network simulations plus an additional 12 attractors. Interestingly, the same 15 perturbations on FVS control nodes that passed the second filtering criterion using the original network were the only combinations of perturbations on FVS control nodes to pass this criterion again in the modified network when using the set of 9 internal-marker nodes. Finally, for these 15 combinations of perturbations on FVS control nodes that passed the second filtering criterion, the steady-state expression value of Dusp6 successfully shifted into the range of the attractors generated from the ESC initial state data.

**Supplementary Data 1:** Excel format. The 68 FVSes in the colorectal cancer signaling network. Each numbered row contains a 13-node FVS in the network.

**Validation of Colorectal Cancer Network and SFA Simulation**

In this study of adaptive resistance to MAPK inhibitor therapy in CRC, the ultimate therapeutic goal is to decrease proliferation and increase apoptosis in tumor cells. In a method adapted from Beal *et al*.[28], we verify that CRC tumors' proliferation and apoptosis signatures are preserved under the SFA simulation of the generic CRC network for individual cell lines or tumors. The underlying concept is that each CRC tumor has specific values for genes related to apoptosis and proliferation based on their normalized expression data and mutational profiles (Supplementary Figure 4a). The genes related to apoptosis and proliferation within the network can be used as signatures of apoptosis and proliferation for the tumor. The corresponding values of these genes are used as a score for the apoptosis and proliferation signatures. We want to verify that after the network is simulated, the steady-state values of the signature genes maintain the original signature scores.

First, we used the Molecular Signatures Database[29] gene sets for "Apoptosis" and "G2M Checkpoint" (proliferation-related genes) to identify 12 genes related to apoptosis and ten genes related to proliferation within the CRC signaling network to be used for the apoptosis and proliferation signatures. Normalized gene expression data for 592 CRC tumors were obtained from The Cancer Genome Atlas[30]. The attractors were estimated with SFA for each tumor sample, initializing the network with the tumor's data profile. Gene Set Variation Analysis[31] was used for each patient tumor to compute apoptosis and proliferation signature scores for the normalized gene expression data and the attractor estimated from the expression and mutational data. Finally, the Spearman Correlation is calculated between the patient data signature score and the attractor signature score for apoptosis and proliferation signatures. The correlation coefficient for the proliferation and apoptosis signature scores was used to evaluate if the network and SFA simulations maintained the appropriate proliferation and apoptotic signatures for each CRC tumor. A strong correlation coefficient indicated that the network and simulations preserved the appropriate signatures and could be used to perform perturbations on FVS control nodes. The signature scores from the patient data and the associated estimated attractors were strongly correlated for apoptosis signature nodes (R=0.76) and proliferation signature nodes (R=0.81) (Supplementary Figure 4b). We concluded that the network and simulation framework could simulate control node perturbations in the system.

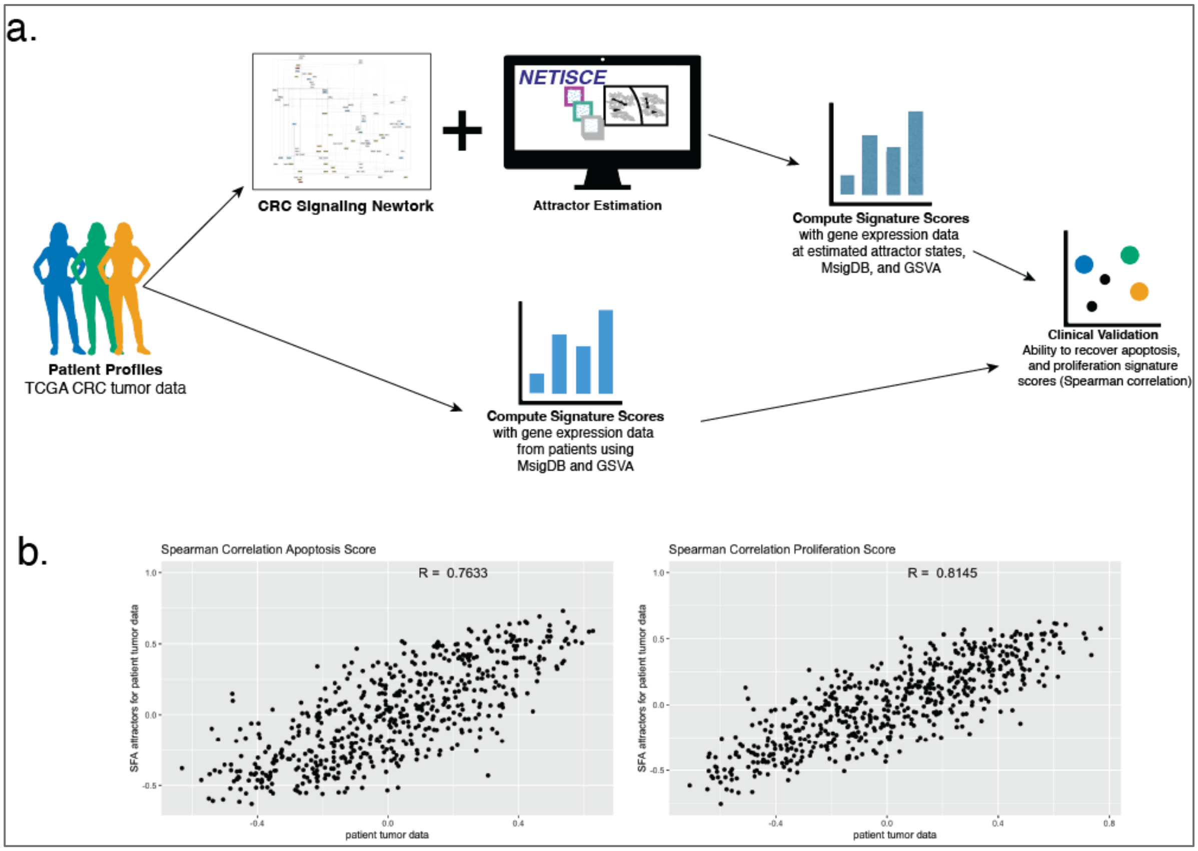


**Supplementary Figure 4: Validation of Colorectal Cancer (CRC) Signaling Network and SFA simulation. a** Overview of the network and simulation validation method, adapted from PROFILE[28]. Using CRC patient tumor data, the underlying concept is that each CRC tumor has specific values for genes related to apoptosis and proliferation based on their normalized expression data and mutational profiles. Our goal is to verify that the proliferation and apoptosis signatures of CRC tumors are preserved under the SFA simulation of the generic CRC network for individual cell lines or tumors. We used the Molecular Signatures Database[29] to identify apoptosis and proliferation-related genes within the CRC signaling network. The corresponding values of these signature genes are used as a score for the apoptosis and proliferation signatures. Starting from patient tumor gene expression data from The Cancer Genome Atlas (TCGA), signature scores for the proliferation and apoptosis-related genes within the network are computed for each patient using Gene Set Variation Analysis[31] (bottom branch). Next, using the network and SFA, an attractor is estimated for each patient, and the apoptosis and proliferation signature scores at the attractor state are computed (top branch). The spearman correlation is calculated between the patient data signature score and the attractor signature score for apoptosis and proliferation signatures. **b** Spearman correlation plots for the apoptosis and proliferation signature scores. The correlation coefficients indicate that the network and simulations preserved the signatures of apoptosis and proliferation.

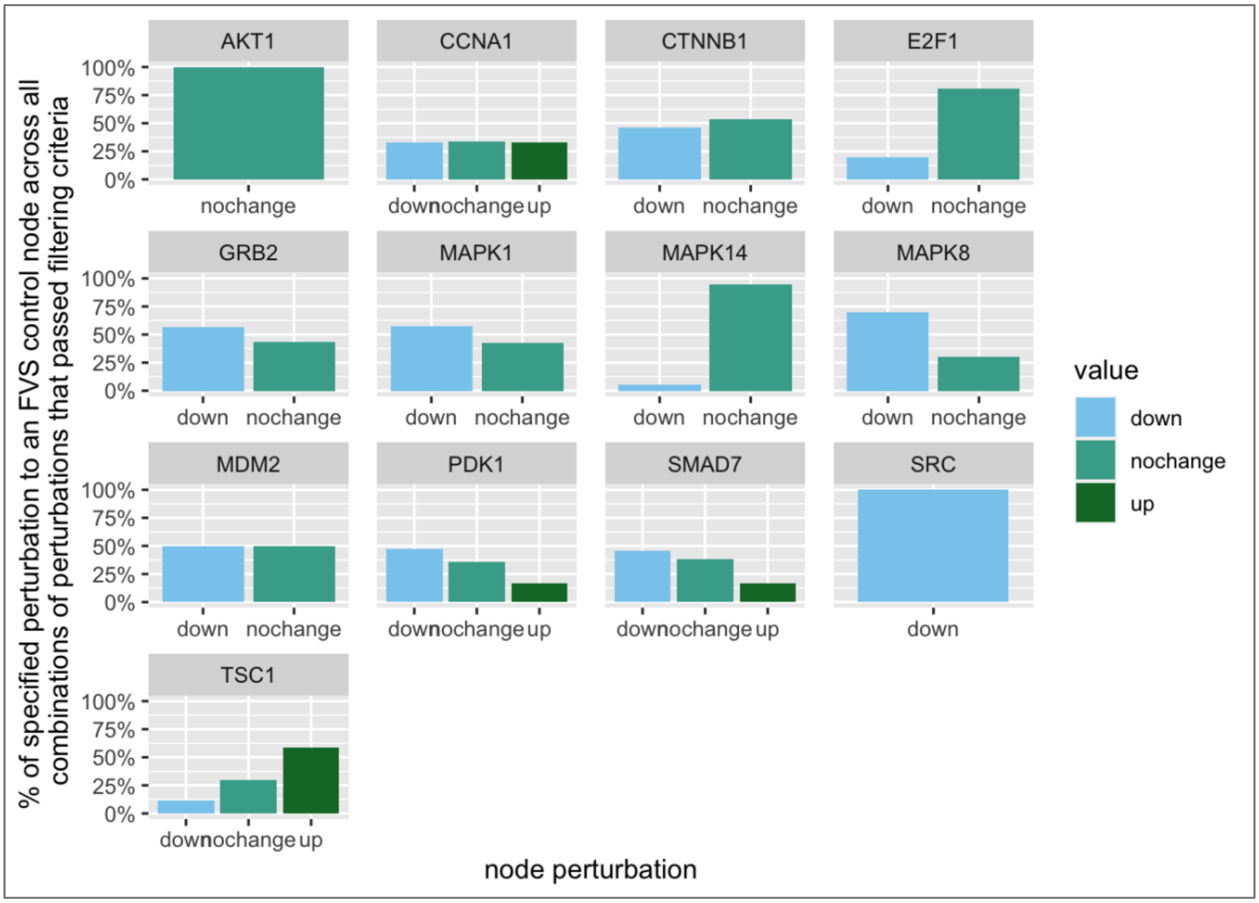


**Supplementary Figure 5:** Bar plots for each FVS control node in the 1,266 perturbations that passed both filtering criteria in the colorectal cancer adaptive resistance reversion simulations when twenty internal-marker nodes are used for Criterion 2. These plots describe the percentages of overexpression and knockout perturbations or no changes to activity for an FVS control node across the 1,266 perturbation sets. The x-axis contains the denoted perturbation (down=knockout, up=overexpression, or no change). The y-axis denotes the % of knockout, overexpression, or no change to that node across all fifteen sets of perturbations on FVS control nodes.

**Feature Importance Analysis in CRC MAPKi Resistance Reprogramming Machine Learning Classification**

We analyzed the top 10 percent of ranked features for the three machine learning classification algorithms (Supplementary Table 7). All three classification methods contain key factors (either FVS control nodes or internal-marker nodes) within their top 10% of ranked features. Additionally, BRAF, DUSP1, and SOS1 are top features in both SVM and Naive Bayes classification, while CTNNB1 and CYCS appear as top features for both Naïve Bayes and Random Forest classifiers. Interestingly, two of the features selected by the Naive Bayes algorithm, DNA_damage, and EGFR, are either source nodes or directly below network source nodes, while the top features for Random Forest and SVM classification appear further downstream in signaling pathways in the network.

**Supplementary Table 7:** Top 10% of ranked features for SVM, Naive Bayes, and Random Forest classification algorithms in CRC MAPKi Resistance Reprogramming simulations.

| **SVM** | **Naïve Bayes** | **Random Forest** |
|---|---|---|
| BRAF | BRAF | MAP2K4 |
| MAP2K1 | EGFR | CYCS |
| MAPK1 | mTOR2 | XIAP |
| RPS6KA1 | FOXO3 | JAK2 |
| SOS1 | CTNNB1 | GRB2 |
| APRY1 | CYCS | ATF2 |
| GAB1 | DIABLO | SOS1 |
| HNF1B | DNA_damage | MAPKAPK2 |
| DUSP1 | DUSP1 | CTNNB1 |
| RAF1 | E2F1 | CDKN1A |

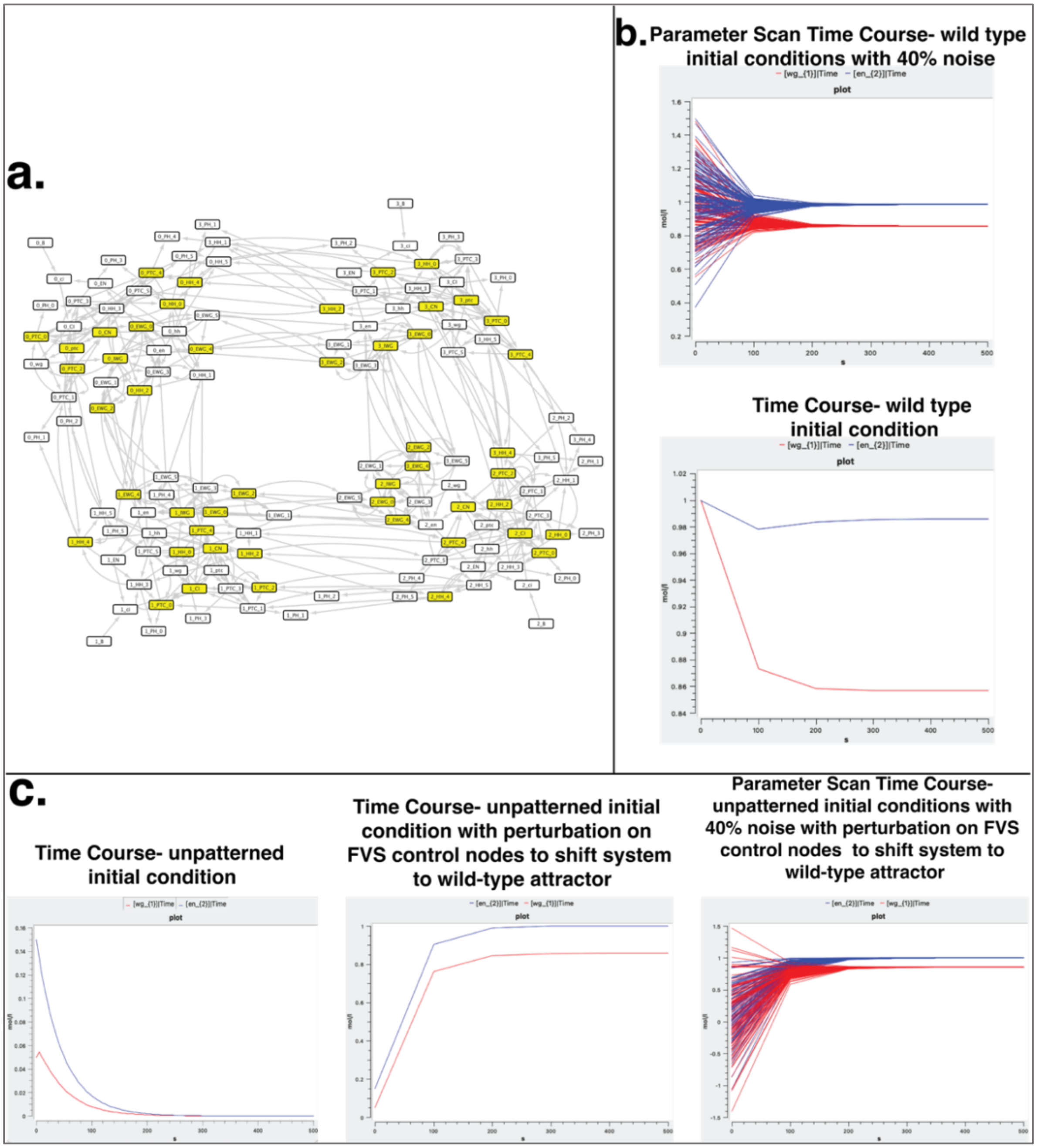


**Supplementary Figure 6: von Dassow Drosophila Segment Polarity Gene ODE model. a** Drosophila Segment Polarity network. The nodes highlighted in yellow comprise the FVS. **b** The lower panel displays the time series simulation using 0% noise for internal-marker nodes wg_1 and en_2. The top panel shows the time course results for the simulations of 1,000 wild-type initial states with 40% noise. For all noisy initial states, the steady-state values of wg_1 and en_2 are the same as those in the simulation with 0% noise. **c** We perform time-course simulations with the unpatterned initial conditions. The first panel shows the time-course results for wg_1 and en_2 under the simulation of the unpatterned initial condition with 0% noise. In the second panel, the results of simulating the unpatterned initial condition with the FVS control show that the steady-state values of the marker nodes shift to the values in the wild-type steady state. The third panel displays the results of simulations of the unpatterned initial conditions with 40% noise with the FVS control applied.

**Supplementary Table 8:** Summary of the NETISCE simulations of control in von Dassow Drosophila Segment Polarity Gene model with noisy initial states. 1,000 sets of three wild-type and three unpatterned initial states with increasing noise levels were provided as initial states for network nodes to NETISCE. The goal of these simulations was for NETISCE to correctly predict that the specified perturbations on FVS control nodes could shift the system from the unpatterned attractor to the wild-type attractor. The second column contains the number of the 1,000 sets per noise level that passed criterion 1. the second column contains the number of the 1,000 sets per noise level that passed criterion 2 and the percentage of the total that passed criterion 2 in parentheses.

| **% noise** | **# passing criterion 1** | **# passing criterion 2** |
|---|---|---|
| 1% | 1000 | 1000 (100%) |
| 5% | 1000 | 996 (99.6%) |
| 10% | 1000 | 987 (98.7%) |
| 20% | 1000 | 969 (96.9%) |
| 30% | 1000 | 926 (92.6%) |
| 40% | 999 | 856 (85.6%) |
| 50% | 965 | 763 (76.3%) |

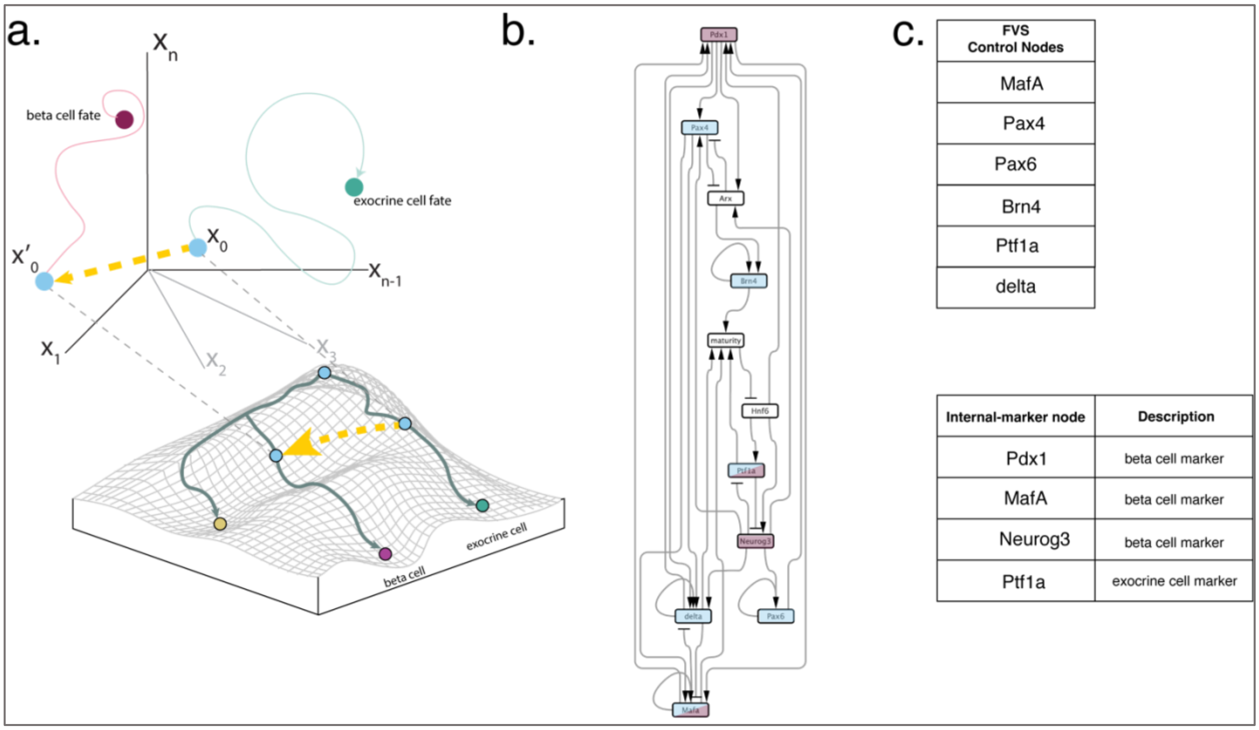


**Supplementary Figure 7: Reprogramming Pancreatic Exocrine Cells to Beta-Cells. a** In this study, we use NETISCE to identify PFVS that shift pancreatic exocrine cells to beta-cells. We initialize the system with gene expression data from exocrine cells ($x_0$) and simulate combinations of perturbations on the FVS control nodes to shift the system to a state (yellow arrow leading to $x'_0$) that leads to beta-cells (magenta circle). **b** Pancreatic cell fate differentiation GRN introduced in Zhou *et al*.[18]. Blue nodes are control nodes, and magenta nodes are internal-marker nodes. **c** Key nodes for the cell fate reprogramming. Presented here is the FVS control node set within the network. Four internal-marker nodes are used to identify targets to reprogram the system towards the beta-cell fate. The phenotypes associated with the internal-marker nodes are denoted in the second column.

**Reprogramming Pancreatic Exocrine Cells to Beta-Cells in NETISCE**

The Zhou *et al.* SDE model of pancreatic cell fate differentiation was used to simulate cell fate reprogramming from pancreatic exocrine cells to beta-cells[18]. After reproducing experimentally-verified exocrine cell reprogramming to beta-cells through MafA, Pdx1, and Ngn3 overexpression, Zhou *et al.* identified that additional Pax4 overexpression and Ptf1a knockout could improve exocrine to beta-cell reprogramming efficiency. Microarray gene expression data for exocrine and beta-cells were obtained from Gene Expression Omnibus (GEO: GSE12025) and normalized using the Voom R package[32]. There were three replicates for each cell type. The pancreatic cell fate differentiation gene regulatory network contains 11 nodes and 34 edges (Supplementary

Figure 7b). Using NETISCE, we estimated the attractors from the exocrine and beta-cell normalized gene expression data and the 100,000 randomly generated initial states. On these 100,006 attractors, we performed k-means clustering. The optimal number of clusters identified by the elbow and silhouette metrics was k=2. One cluster contained the attractors generated from the exocrine cell gene expression. The second cluster included the attractors generated from the beta-cell gene expression.

NETISCE identified one FVS in the network (Supplementary Figure 7c), comprising six nodes: Mafa, Pax4, Pax6, Pou3f4, Ptf1a, and the "delta gene." We simulated 324 combinations of PFVS. Of the 324 combinations of perturbations on FVS control nodes, 86 passed the machine learning classification filtering criterion. We used four internal-marker nodes (Supplementary Figure 7c): Pdx1, Mafa, Neurog (genes upregulated in beta-cells), and Ptf1a (gene upregulated in exocrine cells). The steady-state expression values of the internal-marker nodes in 31 of the 86 attractors calculated from the combinations of perturbations on FVS control nodes were in the range of gene expression values of the beta-cell-associated attractors and thus passed criterion 2 for all three replicates. The FVS contains only a subset of the nodes studied for exocrine cell fate reprogramming in Zhou *et al.:* MafA, Pax4, and Ptf1a. Consistent with their results, the attractor generated from the combination of Ptf1a knockout and MafA overexpression passed both filtering criteria. It is important to note that although the mathematical model of pancreatic cell fate differentiation is stochastic, the deterministic simulations in NETISCE capture the appropriate cell fate reprogramming behaviors.

**Supplementary Table 9:** Summary of the NETISCE simulations in Zhou's Pancreatic Cell Fate Specification model with noisy initial states. 1,000 sets of three exocrine cell and three beta-cell normalized gene expression sets with increasing noise levels were provided as initial states for network nodes to NETISCE. The goal of these simulations was for NETISCE to correctly predict the 31 combinations of perturbations on FVS control nodes that shift the system from the exocrine cell fate to the beta-cell fate. The first column contains, per noise level, the number of the 1,000 sets that passed criterion 1. The second column contains, per noise level, the number (and percentage) of the 1,000 sets that passed criterion 2.

| **% noise** | **# passing criterion 1** | **# passing criterion 2** |
|---|---|---|
| 1% | 1000 | 941 (94.1%) |
| 5% | 1000 | 897 (89.7%) |
| 10% | 1000 | 896 (89.6%) |
| 20% | 1000 | 895 (89.5%) |
| 30% | 1000 | 881 (88.1%) |
| 40% | 1000 | 882 (88.2%) |
| 50% | 1000 | 841 (84.1%) |
